# Multiple Functional Neurosteroid Binding Sites on GABA_A_ Receptors

**DOI:** 10.1101/357574

**Authors:** Zi-Wei Chen, John R. Bracamontes, Melissa M. Budelier, Allison L. Germann, Daniel J. Shin, Krishnan Kathiresan, Ming-Xing Qian, Brad Manion, Wayland W. L. Cheng, David E. Reichert, Gustav Akk, Douglas F. Covey, Alex S. Evers

## Abstract

Neurosteroids are endogenous modulators of neuronal excitability and nervous system development and are being developed as anesthetic agents and treatments for psychiatric diseases. While GABA_A_ receptors are the primary molecular targets of neurosteroid action, the structural details of neurosteroid binding to these proteins remain ill-defined. We synthesized neurosteroid analogue photolabeling reagents in which the photolabeling groups were placed at three positions around the neurosteroid ring structure, enabling identification of binding sites and mapping of neurosteroid orientation within these sites. Using middle-down mass spectrometry, we identified three clusters of photolabeled residues representing three distinct neurosteroid binding sites in the human α_1_β_3_GABA_A_ receptor. Novel intrasubunit binding sites were identified within the transmembrane helical bundles of both the α_1_ and β_3_ subunits, adjacent to the extracellular domains. An intersubunit site in the interface between the β_3_(+) and α_1_(-) subunits of the GABA_A_ receptor pentamer was also identified. Computational docking studies of neurosteroid to the three sites predicted critical residues contributing to neurosteroid interaction with the GABA_A_ receptors. Electrophysiological studies based on these predictions indicate that both the α_1_ intrasubunit and β_3_-α_1_ intersubunit sites are critical for neurosteroid action.

## Introduction

Neurosteroids are cholesterol metabolites produced by neurons^1^ and glia^2^ in the central nervous system (CNS) that are thought to play important roles in both nervous system development and behavioral modulation^3^. Neurosteroid analogues are also being developed as sedative hypnotics^4^, anti-depressants^5^ and anticonvulsants^6^. GABA_a_ receptors, the principal ionotropic inhibitory neurotransmitter receptors in the CNS, have been identified as the primary functional target of neurosteroids. The major endogenous neurosteroids, allopregnanolone and tetrahydroxy-desoxycorticosterone (THDOC), are positive allosteric modulators of GABA_A_ receptors, potentiating the effects of GABA at nanomolar concentrations and directly activating currents at micromolar concentrations. GABA_A_ receptors are members of the pentameric ligand-gated ion channel (pLGIC) superfamily, and are typically composed of two α subunits, two β subunits and one γ or δ subunit^7^. There are 19 homologous GABA_A_ receptor subunits (including six three β and two γ isoforms), with each subunit composed of a large extracellular domain (ECD), a transmembrane domain formed by four membrane spanning helices (TMD1-4), a long intracellular loop between TMD3 and TMD4 and a short extracellular C-terminus. These distinctive structural domains form binding sites for a number of ligands: GABA and benzodiazepines bind to the ECD; picrotoxin to the channel pore^8^; and general anesthetics, such as propofo^9, 10^, etomidate^11^, barbiturates^12^ and neurosteroids to the TMDs^13-18^.

Substantial evidence indicates that neurosteroids produce their effects on GABA_A_ receptors by binding to sites within the TMDs^13-15, 19,20^. Whereas the TMDs of β-subunits are critically important to the actions of propofol and etomidate^11, 21-26^ the α-subunit TMDs appear to be essential for neurosteroid action. Mutagenesis studies in α_1_β_2_γ_2_ GABA_A_ receptors identified several residues in the α_1_ subunit, notably Q^241^ in TMD1, as critical to neurosteroid potentiation of GABA-elicited currents^14,27^. More recent crystallographic studies have shown that in homo-pentameric chimeric receptors in which the TMDs are derived from either α_1_^16^ or α_5_ subunits^17^, the neurosteroids THDOC and pregnanolone bind in a cleft between the α-subunits, with the C3-OH substituent of the steroids interacting directly with.α_1_^Q241^ Neurosteroids are positive allosteric modulators of these chimeric receptors and α_1_^Q241L^and α_1_^Q241W^ mutations eliminate this modulation. These studies posit a single critical binding site for neurosteroids that is conserved across the six α-subunit isoforms^14,27^

A significant body of evidence also suggests that neurosteroid modulation of GABAA receptors may be mediated by multiple sites. Site-directed mutagenesis identified multiple residues that affect neurosteroid action on GABA_A_ receptors, suggestive of two neurosteroid binding sites, with one site mediating potentiation of GABA responses and the other mediating direct activation^14,27^. Single channel electrophysiological studies as well as studies examining neurosteroid modulation of [^35^S]t-butylbicyclophosphorothionate (TBPS) binding, have also identified multiple distinct effects of neurosteroids with various structural analogues producing some or all of these effects, consistent with multiple neurosteroid binding sites^28-30^. Finally, neurosteroid photolabeling studies in the bacterial pLGIC, *Gloeobacter* ligand-gated ion channel (GLIC), demonstrate two neurosteroid binding sites per monomer^31^, one analogous to the canonical intersubunit site and one located in an intrasubunit pocket previously shown to bind propofol^32, 33^ and the inhalational anesthetics^33, 34^ Both of these sites contribute to neurosteroid modulation of GLIC currents, suggesting the possibility of analogous sites in GABA_A_ receptors.

We have developed a suite of neurosteroid analogue photolabeling reagents with photolabeling groups positioned around the neurosteroid ring structure to identify all of the neurosteroid binding sites on GABA_A_ receptors and determine the orientation of neurosteroid binding within each site. Photolabeling was performed in membranes from a mammalian cell line that stably expresses α_1HIS-FLAG_β_3_ receptors (rather than in detergent-solubilized receptors) to ensure that the receptors were in native conformations and environment. Finally, we deployed a middle-down mass spectrometry approach, coupled with a stable-heavy isotope encoded click chemistry tag for neurosteroid-peptide adduct identification to circumvent challenges associated with mass spectrometric identification (predominantly neutral loss) and quantification of neurosteroid-peptide adducts^35^. Using these approaches we have identified three clusters of neurosteroid photolabeled residues on the human α_1_β_3_ GABA_A_ receptor. Computational docking studies, guided by the photolabeling data, were used to describe three binding sites and the orientation of the neurosteroids within each site. The docking studies were also used to predict critical residues to test the contribution of each of these sites to neurosteroid modulation of GABA_A_ currents. Site-directed mutagenesis of these sites and electrophysiological studies indicate that at least two of three structurally distinct sites contribute to allosteric modulation of GABA currents.

## Results

### Development and characterization of allopregnanolone-analogue photolabeling reagents

Allopregnanolone (3α-hydroxy-5α-pregnan-20-one) is a potent, endogenous positive allosteric modulator of GABA_A_ receptors (figure 1a). We synthesized three photolabeling analogues of allopregnanolone in which photolabeling moieties were placed at various positions around the steroid backbone. KK123 has a 6-diazirine photolabeling group on the C5-C6-C7 edge of the sterol, which is a likely binding interface with α-helices^36^ and minimally perturbs neurosteroid structure^37^. KK123 is, however, an aliphatic diazirine and, as such, may preferentially label nucleophilic amino acids^38^. The two other reagents, KK202 and KK200 incorporate a trifluoromethylphenyl-diazirine (TPD) group at either the 3- or 17-carbon. These were designed to sample the space in the plane of the steroid off either the A ring (KK202) or the D ring (KK200). Following UV irradiation, TPD groups generate a carbene which can insert into any bond^39, 40^ Thus, while the TPD groups are bulky and removed several angstroms from the neurosteroid pharmacophore, they should form an adduct precisely at their binding site in the GABA_A_ receptor. Where feasible (KK123, KK202), an alkyne was incorporated in the photolabeling reagents to allow attachment of a fluorophore, purification tag or a mass spectrometry reporter tag (*FLI*-tag) via click chemistry^35^.

**Figure 1.**
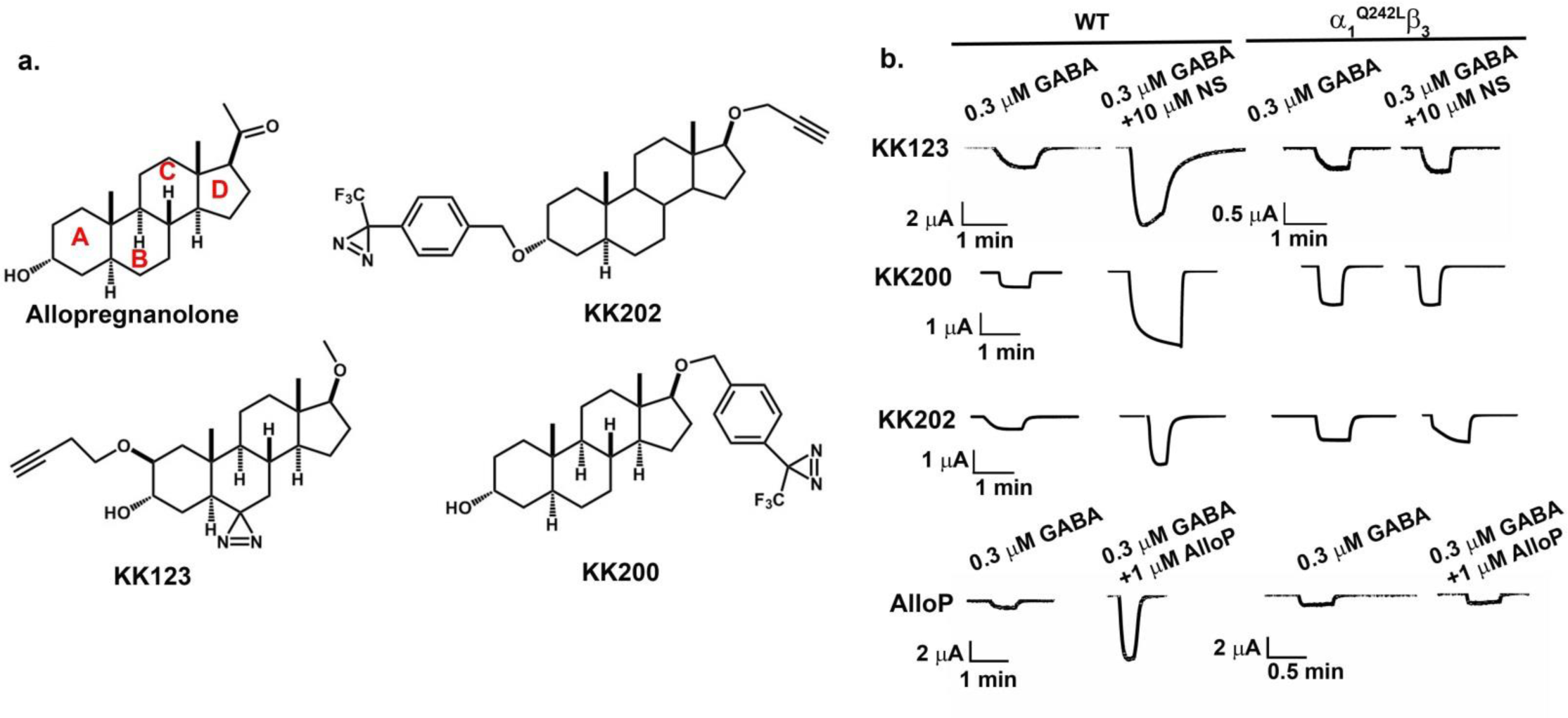
Allopregnanolone analogue photolabeling reagents. a. The structure of allopregnanolone, KK123, KK200 and KK202. b. Allopregnanolone analogue photolabeling reagents potentiate GABA-elicited currents of α_1_β_3_-GABA_A_ receptors; potentiation is blocked by mutation of α_1_^Q242L^β_3_, indicating these photolabeling reagents mimic the action of allopregnanolone.

A useful photoaffinity labeling reagent must bind to the same site on a protein as the ligand it mimics and should produce the same effects on protein functions. To determine if our photoaffinity labeling reagents mimic allopregnanolone as modulators of GABA_A_ receptor function, we assessed modulation of α_1_β_3_GABA_A_ receptors currents in *Xenopus laevis* oocytes, and enhancement of [^3^H]muscimol binding in HEK cell membranes expressing α_1_β_3_ GABA_A_ receptors. KK123 enhanced GABA-elicited (0.3 μM) currents 4.2±3.3 fold at 1 μM (n=5 cells) and 8.2±6.7 fold at 10 μM (n=7). KK123 (10 μM) also directly activated α_1_β_3_ GABA_A_ receptors, eliciting 6.3±3.8% (n=5) of the maximum current elicited by a saturating concentration of GABA. KK123 potentiation of GABA-elicited currents and direct activation were absent in α_1_^Q242L^β_3_ GABA_A_ receptors indicating that KK123 closely mimics the actions of allopregnanolone (the human α_1_ ^Q242L^ mutation is equivalent to rat α_1_^Q242L^ and known to selectively prevent neurosteroid action^14,27^) (figure 1b and table 1). KK200 and KK202 also potentiated GABA-elicited currents at 1 and 10 μM and directly activated the channels at 10 μM (table 1). Positive allosteric modulation by KK202 was somewhat surprising, given that an ether-linked TPD group replaces the 3α-ΟΗ group thought to be critical for neurosteroid action. While the effects of KK200 were abolished in α_1_^Q242L^β_3_receptors, the potentiation by KK202 was reduced by 50% in α_1_^Q242L^β_3_ receptors suggesting that KK202 may have actions at both the canonical neurosteroid site and other binding sites. Since photolabeling experiments were performed in membranes prepared from cells expressing α_1_β_3_ GABA_A_ receptors, we also examined the ability of the photolabeling reagents to enhance [^3^H]muscimol binding in these membranes. A stable HEK-293 cell line was established with tetracycline-inducible expression of human α_1HIS-FLAG_β_3_ GABA_A_ receptors (See Supplemental Methods); receptor density in these membranes was 20-30 pmol [^3^H]muscimol binding/mg membrane protein. The average stoichiometry of the receptors was estimated at two α_1_ subunits and three β_3_ subunits using mass spectrometric (MS) label-free quantitation^41^ (spectral counts). Allopregnanolone enhanced [^3^H]muscimol binding to these recombinant receptors 4-fold with an EC_50_ of 3.9±5.6 μM (Supplemental figure 1). KK123, KK200 and KK202 all enhanced [^3^H]muscimol binding with EC_50_ values similar to or lower than allopregnanolone (supplemental figure 1). Collectively the electrophysiology and radioligand binding data indicate that KK123, KK200 and KK202 are functional mimetics of allopregnanolone.

**Table 1.** Allopregnanolone and the photolabeling analogues modulate and activate α_1_ β_3_-GABA_A_ receptors in *Xenopus laevis* oocytes, tested by two-electrode voltage clamp. Potentiation is expressed as potentiation response ratio, calculated as the ratio of the peak response in the presence of GABA and neurosteroids to the peak response in the presence of GABA alone. Direct activation is expressed as percentage of the response to saturating GABA. Data are shown as mean ± S.D. (number of cells).

| Neurosteroids | Potentiation by 1 μM NS | Potentiation by 10 μM NS | direct activation 10 μM NS |
| --- | --- | --- | --- |
| KK123 | 4.2±3.3 (5) | 8.2±6.7 (7) | 6.3±3.8 (5) |
| KK200 | 1.6±0.2 (5) | 4.9±2.7 (5) | 0.14±0.08 (5) |
| KK202 | 1.4±0.2 (6) | 3.9±2.1 (9) | 8.97±8.9 (6) |
| | Potentiation by 0.1 μM AlloP | Potentiation by 1 μM AlloP | direct activation 10 μM AlloP |
| Allopregnanolone | 4.9±1.3 (7) | 9.7±4.7 (7) | 3.4±1.2 (5) |

To determine whether KK123, which contains an aliphatic diazirine, photolabels GABA_A_ receptors, we utilized the butynyloxy (alkyne) moiety on KK123 to attach a biotin purification tag for selective enrichment of photolabeled GABA_A_ receptor subunits. HEK-293 cell membranes containing α_1_β_3_ GABA_A_ receptors were photolabeled with 15 μΜ KK123, solubilized in SDS and coupled, via Cu^2+^-catalyzed cycloaddition to MQ112 (supplemental figure 2a), a trifunctional linker containing an azide group for cycloaddition, biotin for biotin-streptavidin affinity purification, and a cleavable azobenzene group for elution of photolabeled proteins. The photolabeled-MQ112-tagged receptors were bound to streptavidin beads and eluted by cleavage of the linker with sodium-dithionite. The purified, photolabeled GABA_A_ receptor subunits were assayed by Western blot using anti-α_1_and anti-β_3_. A band at 52-kDa was observed with both α_1_ and β_3_ subunit antibodies in the KK123 photolabeling group (supplemental figure 2b), indicating that both α_1_ and β_3_ subunits are photolabeled by KK123. In control samples photolabeled with ZCM42, an allopregnanolone photolabeling analogue containing a diazirine at the 6-carbon but no alkyne (supplemental figure 2c), neither α_1_ nor β_3_ subunits were purified. These data indicated that KK123 can photolabel both α_1_ and β_3_ subunits and is thus an appropriate reagent to use for site-identification.

### Identification of KK123 photolabeling sites on α_1_ and β_3_ subunits of GABA_A_ receptors

Identification of sterol adducts in hydrophobic peptides has been impeded by multiple challenges including peptide insolubility during sample digestion, ineffective chromatographic separation of hydrophobic TMD peptides and neutral loss of sterol adducts from small hydrophobic peptides during ionization and fragmentation. To circumvent these problems we employed middle-down mass spectrometry to analyze GABA_A_ receptor TMD peptides and their sterol adducts. This approach identifies each TMD as a single, large peptide and attenuates neutral loss of adduct, facilitating identification of the sites of neurosteroid incorporation. In our studies, α_1HIS-FLAG_β_3_ GABA_A_ receptors were photolabeled in native HEK cell membranes. The photolabeled proteins were then solubilized in n-dodecyl-β-D-maltoside (DDM)-containing lysis buffer. The pentameric GABA_A_ receptors were purified using anti-FLAG agarose beads and eluted receptors were digested with trypsin in the presence of MS-compatible detergent DDM. These conditions generated peptides containing each of the GABAA receptor TMDs in their entirety. The peptides were separated using PLRP-S nano-liquid chromatography and analyzed on a Thermo Elite orbitrap mass spectrometer. This work flow (see supplemental figure 3) minimized protein/peptide aggregation, simplified MS-1 level identification of TMD-sterol adducts and optimized fragmentation of TMD peptides and their adducts. All eight of the TMD peptides were reliably sequenced with 100% residue-level coverage. In addition, the covalent addition of neurosteroid to the TMD peptides increased the hydrophobicity of TMD peptides and shifted their chromatographic elution to later retention times (supplemental figure 3). The delayed retention time was used as a critical criterion for identification of photolabeled peptides.

Two photolabeled peptides were found in the mass spectra of tryptic digests of α_1_β_3_ GABA_A_ receptors photolabeled with KK123. A KK123 adduct of the a1-TM4 peptide, ^398^IAFPLLFGIFNLVYWAT<u>Y^KK123^</u>LNREPQLK^423^ (m/z= 875.503, z=4), was identified (add weight of KK123 = 316.27). Site-defining ions in the fragmentation spectra identified the site of KK123 insertion as Y^415^, at the C-terminus of a_r_TM4 (underlined in the sequence above; see figure 2a). In a separate series of experiments, α_1_β_3_ receptors were photolabeled with KK123 which was then coupled to *FLI*-tag using click chemistry. *FLI*-tag, an azide-containing tag, adds both charge and a heavy/light stable isotope pair to a photolabeled peptide, enhancing identification by creating doublets in the MS1 spectra^35^. MS1 level search for pairs of ions differing by 10.07 mass units found two peptide ion features (m/z=1073.246 and m/z=1076.580, z=3) that had identical chromatographic retention times (figure 2b). Fragmentation spectra revealed both of these peptides as β_3_-TM4 peptide (^429^IVFPFTFSLFNLVYW<u>Y^KK123^</u>YVN^445^) with a KK123-FLI-tag adduct (adduct mass=672.432 and mass=682.441) on Y^442^(figure 2c). In the fragmentation spectrum, ions containing KK123 + light *FLI-tag* (figure 2c, black) were different by 10.07 mass units than the corresponding fragment ions from KK123 plus heavy *FLI-* tag (figure 2c, red), confirming that KK123 photolabels Y^442^ of the β_3_ subunit. β_3_-Y^442^is located on the C-terminus of β_3_-TM4 in a homologous position to α_1_-Y^415^, the KK123 photolabeling site in α_1_-TM4 (figure 2d, upper right panel). Thus, KK123 labeling data identified two discrete sites, one in α_1_ and the other in β_3_. We employed additional photolabeling reagents containing TPD groups arrayed around the sterol backbone to confirm whether the KK123-labeled residues represent neurosteroid binding sites and to determine the orientation of the neurosteroids in these sites.

**Figure 2.**
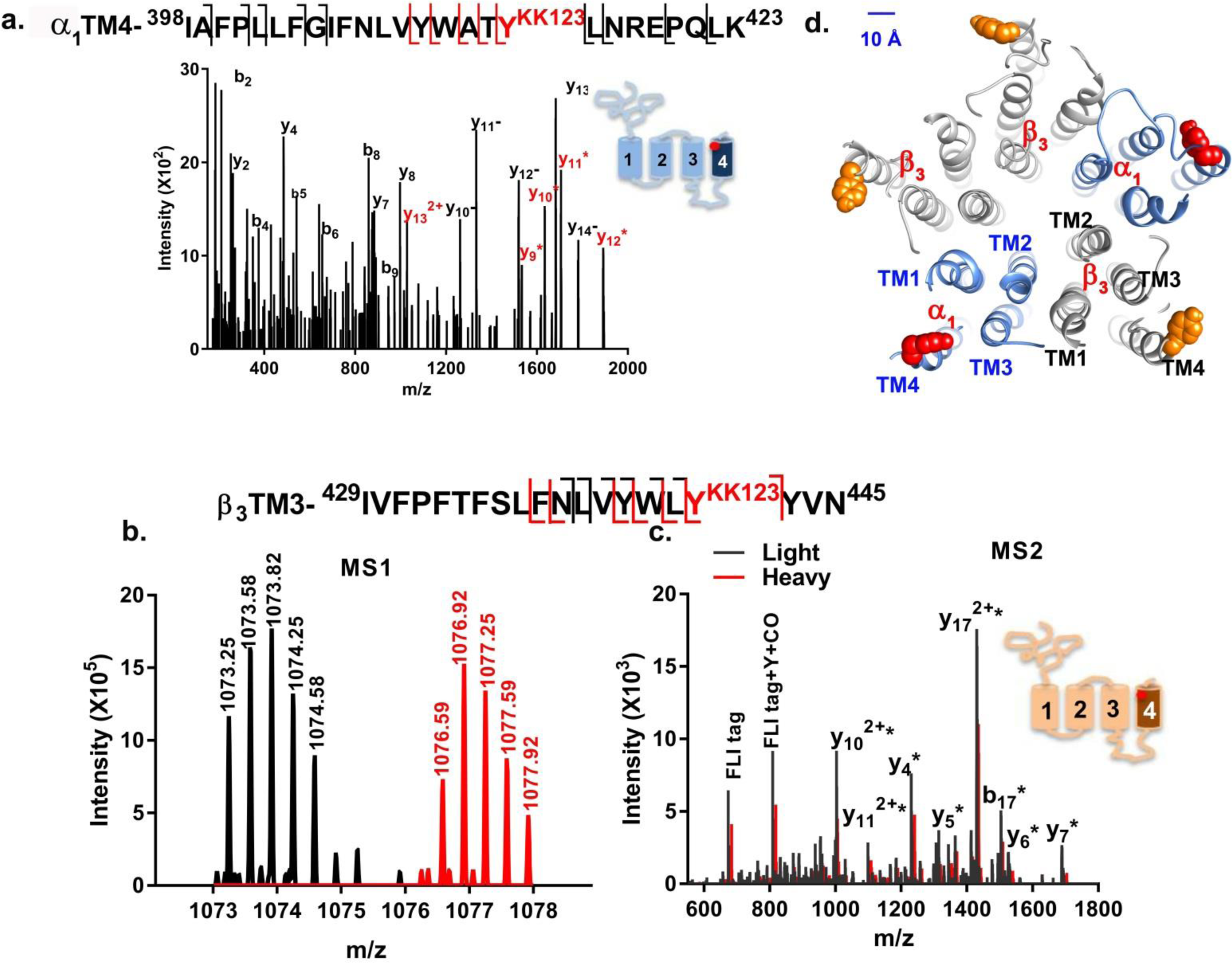
KK123 photolabels α1-TM4 and β3-ΤΜ4 peptides. a. MS fragmentation spectrum of a KK123 photolabeled a_1_-TM4 peptide (m/z=875.503, z=4). The y ions containing KK123 adduct are colored red. Fragment ions y_10_^−^ to y_14_^−^ represent neutral loss of the KK123 adduct. b. MS1 pair of light and heavy form of *FLI* tag-KK123 photolabeled β_3_ TM4 peptide (m/z=1073.246 and m/z=1076.580, z=3). c. An overlay of light (black) and heavy (red) MS fragmentation spectra of *FLI* tag-KK123 photolabeled β_3_ TM4 peptide. The KK123 adduct-containing b or y ions are labeled with *. d. KK123 photolabeled residues are shown in an homology model of the structure of an α_1_β_3_ GABA_A_ receptor. In the α_1_ subunit the lableled TM4 tyrosine points toward TM1, whereas in the β_3_ subunit, the labeled tyrosine residue points toward TM3.

### Photolabeling sites identified by KK200 and KK202

KK200, which has a TPD photolabeling group attached at C17 on the steroid backbone has been previously used to map neurosteroid binding sites on GLIC^31^. Analysis of α_1_β_3_receptors photolabeled with 15 μM KK200 detected two photolabeled TMD peptides: An α_1_-TM4 peptide, ^398^IAFPLLFGIF<u>N^KK200^</u>LVYWATYLNREPQLK^423^, was photolabeled with KK200 (m/z=898.002, z=4); site defining ions in the fragmentation spectra identified N^408^as the modified residue (figure 3a). The N^408^ residue (N^407^ in rat) has previously been shown to be critical to neurosteroid potentiation of GABA-elicited current^s14 15^ A β_3_-TM3 peptide, ^280^AIDMYLMGC^nem+dtt^FVFVFLALLEYAFVNYIFF<u>GR^KK200^</u>GPQR^313^ (m/z=1188.352, z=4; N-ethylmaleimide, NEM; 1, 4-dithiothreitol, DTT; alkylation adduct), was also photolabeled with KK200. Fragmentation spectra narrowed the possible sites of adduction to G^308^or R^309^, both at the junction of TM3 with the M3-M4 intracellular loop (figure 3b).

**Figure 3.**
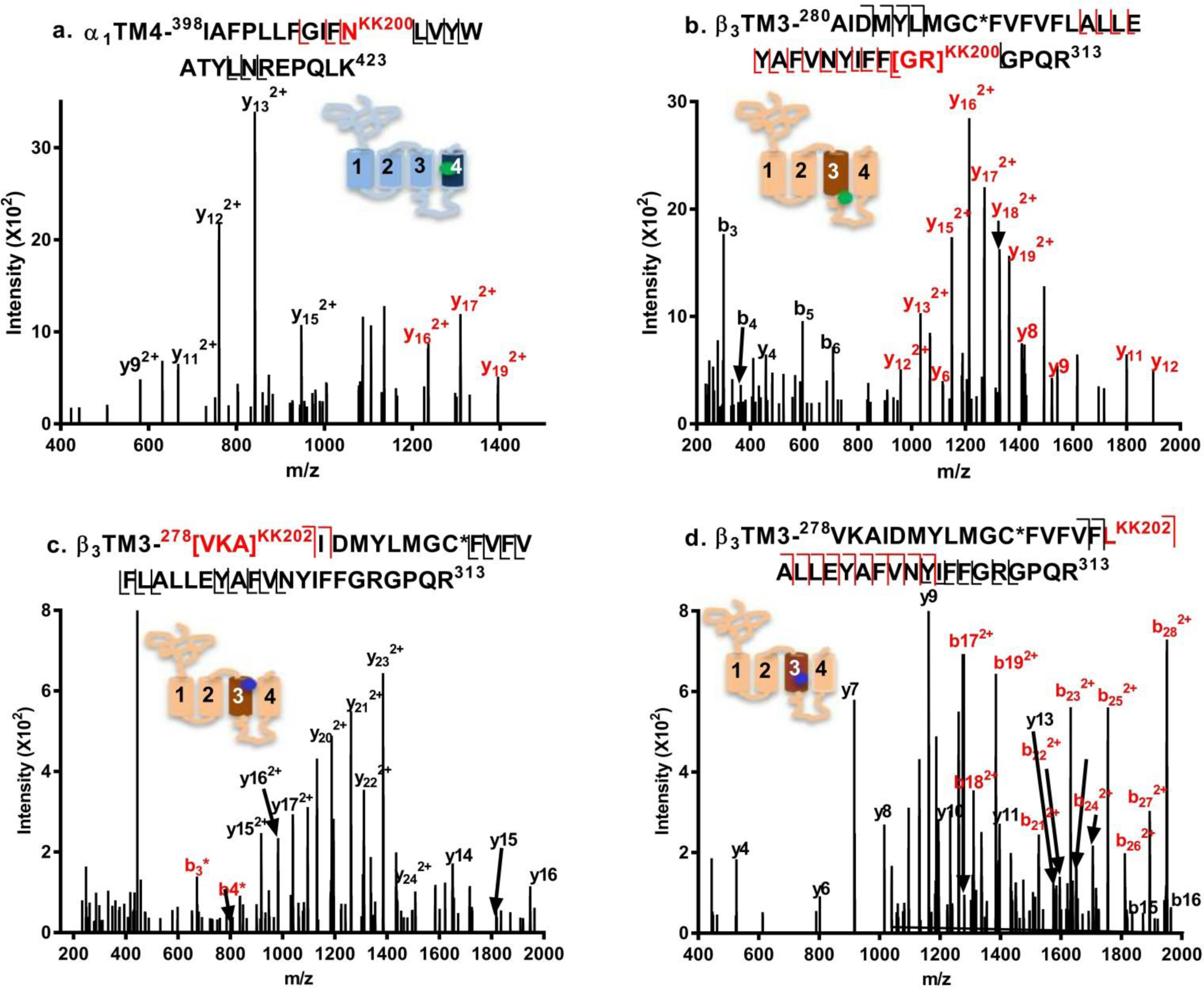
Fragmentation spectra of KK200- and KK202-photolabeled GABA_A_ receptor peptides. a. An a_1_TM4 peptide (m/z=898.002, z=4) is photolabeled by KK200 at N^408^; b. Α β^3^ TM3 peptide (m/z=1188.352, z=4) is photolabeled by KK200 at [^308^GR^309^] c and d. β_3_ TM3 peptides (m/z=811.453, z=6) are photolabeled by KK202 at ^226^VKA^228^ (c) and L^294^ (d). Fragment ions labeled in red contain the neurosteroid adduct.

Analysis of GABA_A_ receptors photolabeled with KK202 (figure 3 c and d), identified two photolabeled peptides eluting two minutes apart. Both peptides were identified as the β_3_-ΤΜ3 peptide, ^278^<u>VKA</u>IDMYLMGC^nem^FVFVF**<u>L</u>**ALLEY AFVNYIFFGRGPQR^313^ (m/z=811.453, z=6). Fragmentation spectra of the earlier eluting peptide localized labeling to a 3 residue sequence, ^278^VKA^280^, at the N-terminus of β_3_-Μ3 (figure 3c). The fragmentation spectrum of the later eluting peptide, identified L^294^ as the site of adduction (figure 3d). (n.b. The different retention time of the two photolabeled peptides is likely due to differences in peptide conformation and surface hydrophobicity resulting from incorporation of the photolabeling reagent into different residues.)

### Allopregnanolone prevents photolabeling by neurosteroid analogue photolabeling reagents

An important test of whether the photolabeled sites constitute specific allopregnanolone binding sites is the ability of excess allopregnanolone to competitively prevent photolabeling. Photolabeling studies for site identification were performed using 15 μΜ photolabeling reagent and achieved levels of labeling efficiency varying from 0.06-3.0 % (supplemental table 1). Since allopregnanolone has limited aqueous solubility (about 30 μΜ) and a large competitor excess is needed to demonstrate competition (particularly with an irreversibly bound ligand), we were limited to studying competition at the photolabeled residues that could be detected following photolabeling at a concentration of 3 μΜ. Accordingly, we measured the photolabeling efficiency obtained following photolabeling of α_1_β_3_ GABA_A_ receptors with 3 μM KK123, KK200 or KK202 in the presence or absence of 30 μΜ allopregnanolone. KK123 photolabeled both α_1_ Y^415^ (0.5% efficiency) and β_3_ −Y^442^ (0.3% efficiency). For both of these residues, photolabeling was reduced by > 90% in the presence of excess allopregnanolone (figure 4a). KK200 photolabeled β_3_ –G^238^/R^309^ (0.18% efficiency) and labeling was reduced by 98% in the presence of allopregnanolone. KK202 labeled both β_3_-L^294^ (0.34 % efficiency) and β_3_-^278^VKA^280^ (0.21 % efficiency) in TM3; labeling of both of these sites was undetectable in the presence of 30 μM allopregnanolone. Studies were also performed to determine if the allosteric agonist GABA (1 mM) enhanced photolabeling by 3 uM KK123 or KK200. Labeling efficiency was not significantly enhanced in the presence of GABA. This suggests that there is a small difference in neurosteroid affinity for closed vs. open/desensitized states which is consistent with the fact that neurosteroids have very low efficacy as direct activators of GABA_A_ receptors^42^

**Figure 4.**
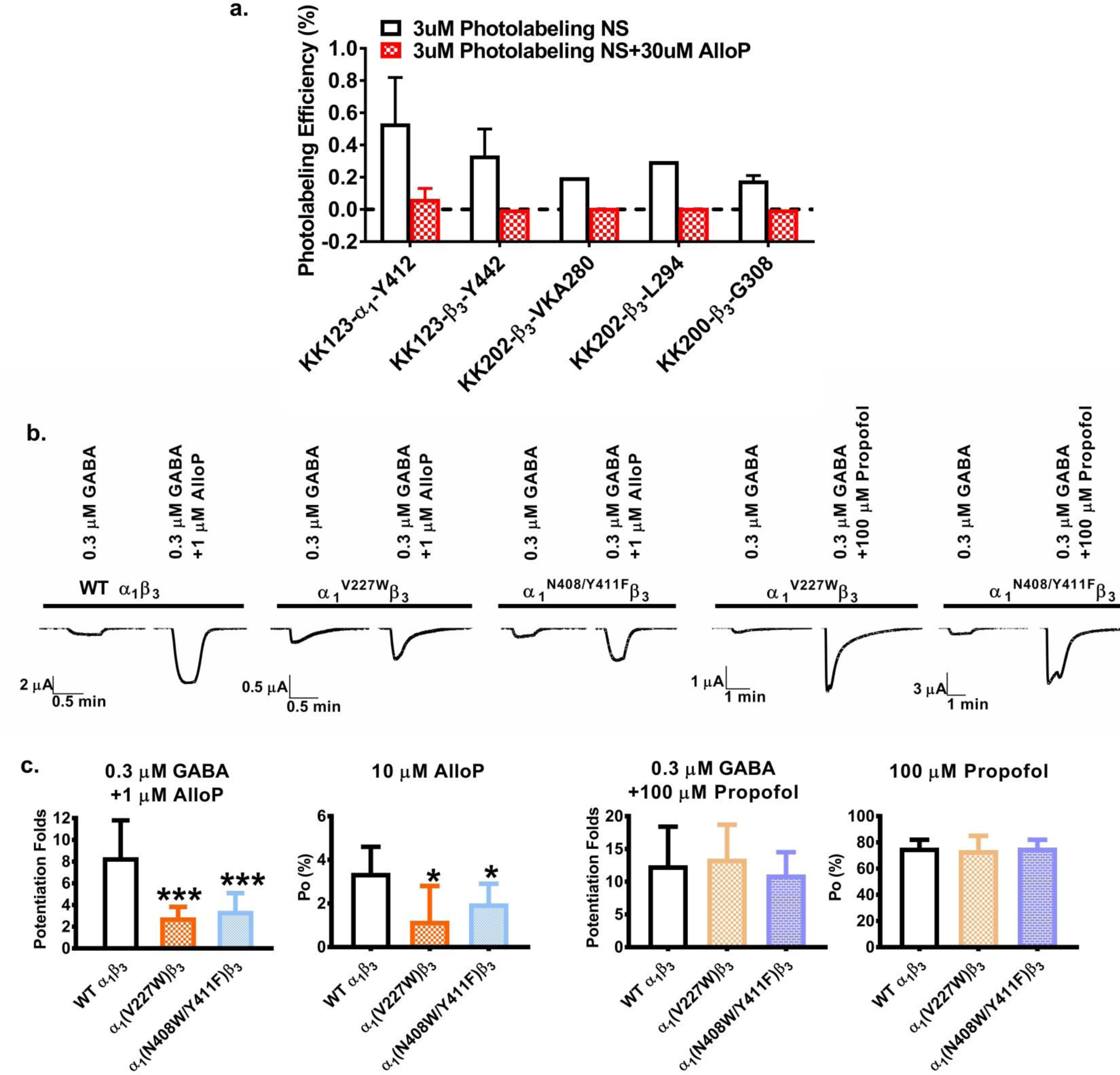
The α_1_ subunit is critical for neurosteroid modulation of α_1_β_3_ GABA_A_ receptors. a. Photolabeling with 3 μM KK123, KK200, and KK202 is prevented by labeling in the presence of 30 μM allopregnanolone; b. Current traces of allopregnanolone and propofol effects on wild-type α_1_β_3_, α_1_^V227W^β_3_ and α_1_^N408W/Y114F^ GABA_A_ receptors. c. Potentiation of GABA-elicited currents and direct activation of α_1_β_3_ GABA_A_ receptors by allopregnanolone is reduced in α_1_^V227W^β_3_ receptors. The α_1_^V227W^β_3_ mutation has no effect on propofol action.

### Structural characterization of the photolabeling sites

The six residues photolabeled by KK123, KK200 and KK202 were examined in a model of the α_1_β_3_ receptor created by threading the aligned sequence of the α_1_ subunit on the structure of the β_3_ subunit (PDB 4COF)^43^. The photolabeling sites grouped into 3 clusters: cluster 1 (blue circle)-β_3_-L^294^ (KK202) and β_3_-G^308^/R^309^ (KK200); cluster 2 (red circle)-α_1_Y^415^ (KK123) and α_1_-N^408^ (KK200) and; cluster 3 (brown circle)-β_3_-Y^442^(KK123) and β_3_-^278^VKA^280^ (KK202) (figure 5a).

**Figure 5.**
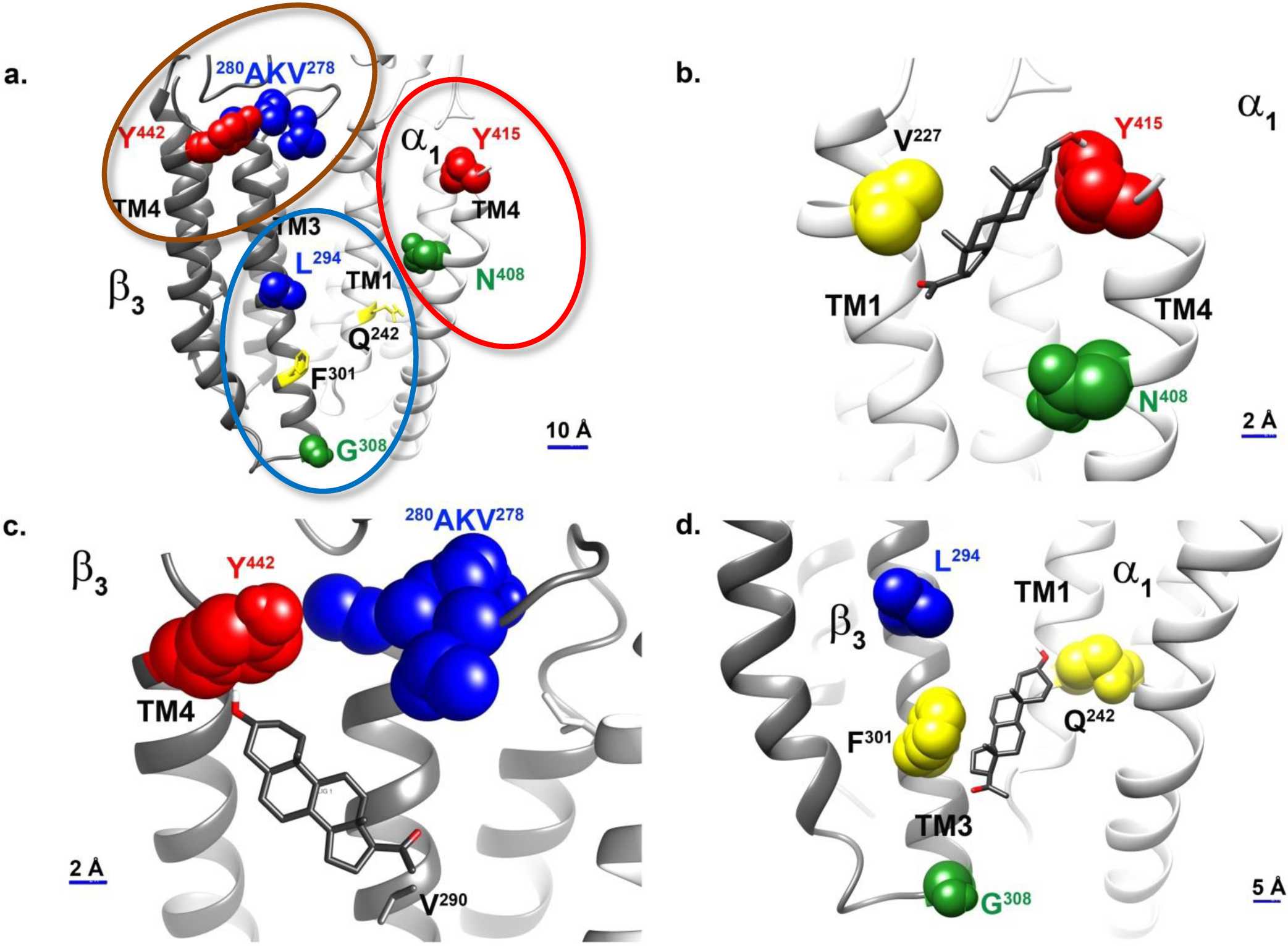
Allopregnanolone docking in the three neurosteroid binding sites identified by photolabeling. a. The six photolabeling sites identified by photolabeling with KK123, KK200, and KK202 grouped into 3 clusters: β_3_(+)/α_1_(-) intersubunit sites (blue circle), β_3_ intrasubunit sites (brown circle), and α_1_ intrasubunit sites (red circle); Docking of allopregnanolone to the α_1_ intrasubunit sites (b), β_3_ intrasubunit sites (c), and β_3_ (+)/α_1_ (-) intersubunit sites (d). Residues photolabeled by KK123, α_1_-Y^415^ and β_3_-Y^442^, are colored red; Residues photolabeled by KK200 α_1_-N^408^ and β_3_-G^308^, are colored green; and residues photolabeled by KK202 β_3_-^278^VKA^280^ and L^294^ are colored blue. Residues previously identified as contributing to an intersubunit neurosteroid binding site, α_1_-Q^242^, and β_3_-F^301^ are shown in yellow, as is α_1_-V^227^ a residue in the α_1_ intrasubunit site shown to affect neurosteroid action by site-directed mutagenesis.

In cluster 1 (blue circle, figure 5a), β_3_-L^294^faces into the β(+)/α(-) intersubunit cleft and G^308^/R^309^is at the junction between the bottom of TM3 and the TM3-4 intracellular loop. G^308^/R^309^ is 2 a-helical turns below β_3_-F^301^(i.e. towards the intracellular terminus of TM3), a site previously photolabeled by 6-azi pregnanolone in β_3_ homomeric receptors^13^. These data support neurosteroid binding in the β(+)/α(-) interface, consistent with the canonical THDOC and pregnanolone binding sites identified in crystal structures of α_1_(+)/α_1_(-) interfaces in chimeric proteins^16^ and in SCAMP studies of α_1_β_2_γ_2_receptors^18^. The pattern of labeling also indicates that the A ring of the steroid is oriented upwards in the intersubunit cleft toward the center of the membrane, the D-ring is pointing toward the intracellular termini of the TMDs and the C5-C6-C7 edge of the steroid is pointing toward the β_3_(+) side of the cleft; this is consistent with the orientation of THDOC determined in the X-ray structure of α_1_-GLIC^16^. Cluster 1 corresponds to a β_3_(+)/α_1_ (-) intersubunit site.

In cluster 2 (red circle, figure 5a), N^408^ and Y^415^ are both on the C-terminal end of α_1_-TM4, facing toward TM1 within the same α_1_ subunit, consistent with an α_1_ intrasubunit neurosteroid binding site. N^408^, the residue labeled by the C17-TPD of KK200 is two α-helical turns closer to the center of TM4 than is Y^415^, the residue labeled by the C6-diazirine of KK123. This labeling pattern suggests that neurosteroids orient in this site with the A-ring pointing toward the extracellular domain and the D-ring facing to the center of the TMD. Cluster 2 corresponds to an α_1_ intrasubunit site.

In cluster 3 (brown circle, figure 5 a), Y^442^ is located at the C-terminal end of β_3_-ΤΜ4 and ^278^VKA^280^ is located on the TM2-TM3 loop near the extracellular end of β_3_-ΤΜ3. The adjacency of these two photolabeling sites suggests an intrasubunit neurosteroid binding site at the extracellular end of β_3_, analogous to the α_1_ intrasubunit site. The labeling of ^278^VKA^280^ in the extracellular loop by the C3-TPD group of KK202 suggests that neurosteroids orient in this site with the A-ring facing the extracellular domain. Cluster 3 corresponds to a β_3_ intrasubunit site.

### Molecular dynamic simulations and docking of neurosteroids

An homology model of the α_1_β_3_ GABA_A_ receptor based on the structure of a β_3_ homomeric GABA_A_ receptor (PDB 4COF)^43^ was used to examine the preferred energetic poses of neurosteroid binding to the three binding sites. The homology model was embedded in a POPC bilayer and the structure refined by molecular dynamics. We then docked each of the three photoaffinity labeling reagents as well as allopregnanolone to each of the proposed binding sites, using a time course series of snapshots from the simulation trajectory to account for receptor flexibility. All of the neurosteroid photolabeling reagents docked in the three sites; the identified sites are relatively shallow with respect to the protein-lipid interface. Moreover, the neurosteroid analogues were all found to adopt multiple poses in each of the sites with minimal energy differences between the poses (see supplementary Materials and Methods). Photolabeling data combined with the docking scores (binding energy) and population of a given pose were used to guide selection of the preferred steroid orientation in each site.

In the α_1_ intrasubunit site, the poses clustered between TM1 and TM4. The preferred pose (figure 5b) for allopregnanolone (lowest energy cluster of poses) shows the A-ring oriented toward the ECD with the walls of the predicted binding site lined on one side by N^408^ and Y^415^ and on the other by V^227^. Docking of KK200 in this site has a similar orientation with the A-ring oriented toward the ECD and the TPD group on the D-ring proximal to N^408^ (supplemental figure 5a). Docking of KK-123 shows a preferred pose in which the A-ring is oriented towards the ECD and the C6-diazirine proximal to Y^415^ (supplemental figure 5b). These data elucidate a prior finding that mutations to N^408^ and Y^411^ eliminate potentiation by steroid analogues that lack a hydrogen bonding group on the D-ring^44^.

In the β_3_ intrasubunit site the poses are clustered between TM3 and TM4. Allopregnanolone preferred a pose with the A-ring oriented towards the ECD near Y^442^ and the D-ring proximal to V^290^ (figure 5c). KK123 was found to dock at the top of the TM helices with the A ring oriented towards the ECD, placing the 6-diazirine in proximity to Y^442^ (supplemental figure 5c). KK202 was found to dock in a similar orientation but lower in the TM region, with the TPD group in proximity to A^280^ and Y^442^ (supplemental figure 5d).

At the intersubunit site the preferred pose for allopregnanolone was one of the lowest energy clusters of poses with the A ring proximal to α_1_-Q^242^ (equivalent to rat α_1_-Q^241^), the D ring pointing toward the cytoplasmic termini of the TMDs and the D ring facing β_3_-F^301^ (figure 5d). Docking of KK200 showed a similar orientation although shifted slightly upwards towards the ECD, placing the A ring near α_1_-Q^242^, benzene ring of the TPD near β_3_-F^301^ and the diazirine in proximity to G^308^ (supplemental figure 5e). The preferred pose of KK202 was closer to TM3 of the β_3_ subunit with the A ring near α_1_-Q^241^ and the D ring near the β_3_-F^301^ placing the diazirine in proximity to β_3_-L^294^ (supplemental figure 5f).

As a confirmation of our docking, we also docked allopregnanolone to the β_3_ homomeric receptor (PDB 4COF^43^) and to the homomeric β_3_-α_3_ chimera in which all of the TMDs are from α_5_ (PDB 5CO8F^17^). Allopregnanolone docked to the intrasubunit sites in β_3_ and α_3_ with the same preferred poses that we selected in our homology model based on our docking and photolabeling criteria.

### Effects of mutations in putative binding sites on neurosteroid action

The β_3_-α_1_ intersubunit binding site identified in our photolabeling studies has been extensively validated by site-directed mutagenesis as a functionally important site. Mutations on the α(-) side of the interface, including Q^241(rat)^L/W and W^246(rat)^L have been shown to eliminate neurosteroid potentiation and gating of α_1_β_2_γ_2_ GABA_A_ receptors^14,27^ Mutations on the β(+) side of the interface, including F^301^ A and L^297^ A have also been shown to partially reduce neurosteroid effect^17^. In the current study we showed that α_1_^Q242L^β_3_ prevented the action of allopregnanolone, KK123 and KK200, while reducing the effect of KK202 in α_1_β_3_ receptors, confirming the β-α interface as a functionally significant neurosteroid binding site and validating the relevance of our photolabeling reagents (figure 1b).

Based on computational simulation and docking results, we also identified residues in the proposed α_1_- and β_3_ intrasubunit binding sites that we predicted could be involved in allopregnanolone binding or action (see supplemental tables 2 and 3 for all mutated subunits tested). N^408^ and Y^411^ in α_1_-TM4 line one side of the putative α_1_ intrasubunit site and V^227^ in α_1_-TM1 lines the other (figure 4b).α_1_^N407(rat)A^and α_1_^Y410(rat)W^ mutations have previously been shown to prevent neurosteroid potentiation of GABA-elicited currents in α_1_β_2_γ_2_ GABA_A_ receptors_15_. Our data confirm that the double mutant α_1_^N408W^/^Y411F^β_3_ reduces allopregnanolone potentiation of GABA-elicited currents and eliminates allopregnanolone direct activation in α_1_β_3_ receptors (p < 0.01 and *p<0.05 vs α_1_β_3_wild-type). Allopregnanolone (1 μM) potentiation of GABA-elicited currents and direct activation (10 μM) of α_1_^V227W^ β_3_ receptors was also reduced by 65% in comparison to α_1_β_3_ wild-type (***p<0.05; figure 4b and c). To test whether these mutations selectively affected neurosteroid actions, we also compared the effects of propofol in α_1_^N408W/Y411F^β_3_ to its effect on wild-type α_1_β_3_ receptors. Propofol action was not affected, indicating a selective effect on neurosteroid action (figure 4b and c). The finding that multiple mutations lining the α_1_-intrasubunit binding pocket selectively reduce allopregnanolone action buttresses the evidence that the photolabeled residues identify a specific, functionally important neurosteroid binding site.

Multiple mutations within the putative β_3_-intrasubunit binding site were also tested. However, none of the mutations significantly altered potentiation or activation by allopregrenanolone (See supplemental table 2 and 3 for all of the mutations that were tested). These data suggest that allopregnanolone occupancy of the β_3_ intrasubunit site does not contribute to channel gating. Interestingly, direct activation of α_1_β_2_^Y284F^γ_2_receptors by THDOC has previously been shown to be markedly reduced in comparison to wild type receptors^15^, although we found no significant effect of the β_3_-Y^284^ mutation in α_1_β_3_receptors (supplemental tables 2 and 3). The difference in results between experiments in α_1_β_3_ and α_1_β_2_γ_2_ GABA_A_ receptors suggests possible isoform specificity in the functional effects of neurosteroid binding at a β-intrasubunit site.

## Discussion

Collectively the photolabeling, modeling and functional data indicate that heteropentameric α_1_β_3_ GABA_A_ receptors contain at least 7 binding sites for neurosteroids, of 3 different types. The use of multiple photolabeling reagents also enabled determination of the orientation of neurosteroids in each proposed class of sites. At least 2 of these classes are involved in producing the allosteric effect of steroids, the β_3_-α_1_ intersubunit site (2 copies per receptor) and the α_1_ intrasubunit site (2 copies). Mutations of residues in the proposed β_3_ intrasubunit site (3 copies) had no effect on modulation by allopregnanolone although residues were labeled by two photolabeling reagents and labeling was prevented by excess allopregnanolone. Accordingly the functional significance of this proposed site is not known.

Previous, site-directed mutagenesis studies using electrophysiology readout identified multiple residues, including α_1_-Q^241^, N^407^, Y^410^, T^236^ and β_3_-Υ^284^, that selectively contribute to the positive allosteric effects of neurosteroid^s14, 27^. Based on homology to the structure of the muscle nicotinic acetylcholine receptor^45^, it was hypothesized that there are two neurosteroid binding sites on GABA_A_ receptors: an α_1_-intrasubunit site spanning Q^241^ and N^407^ and an intersubunit site between β_3_-Υ^284^ and α_1_-T^236^. Subsequent data^16-18^ have clearly established the existence of a β-α intersubunit site. Our photolabeling experiments and homology modeling based on the only GABA_A_ receptor structure^43^, now show that the previously identified residues contribute to multiple distinct neurosteroid binding sites, albeit differently than originally proposed. It is noteworthy that the α_1_-intrasubunit site was not identified in the X-ray crystallographic structures of α_1_-GLIC chimeras bound with THDOC or the α_5_-β_3_ chimera bound with pregnanolone. This is likely because the proteins with steroid bound in the intrasubunit site did not form stable crystals.

Interestingly, mutations in either the β_3_-α_1_ intersubunit site or the α_1_-intrasubunit site can ablate both potentiation and direct activation by allopregnanalone, indicating that there are not distinct sites mediating potentiation and direct activation; rather both sites contribute free energy that can enhance gating of the channel. However, the observation that mutations in either binding site can largely eliminate neurosteroid effect indicates that these two sites do not function completely independently.

In light of the demonstration of multiple neurosteroid binding sites in α_1_β_3_ GABA_A_ receptors, the possibility of additional isoform specific sites must be considered. The strong sequence homology between the TMDs of the six α-subunits and three β-subunits suggests that there will not be large isoform differences in the intersubunit site^27^. In contrast, the contribution of ECD residues to the α- and β-intrasubunit sites suggests possible isoform specific differences. While there are no available structures for the γ and δ subunits, the sequence homology between these subunits and α and β subunits suggests that there may also be intrasubunit neurosteroid binding sites in these isoforms. The existence of multiple sites in which neurosteroids bind with different orientation may also offer some explanation for the difficulty in identifying neurosteroid antagonists^46^ and for the differences in single channel electrophysiological effects of various neurosteroid analogue^s28, 30^. The possibility of multiple isoform-specific sites with distinct patterns of neurosteroid affinity, binding orientation and effect offers the exciting potential for the development of isoform-specific agonists, partial agonists and antagonists with targeted therapeutic effects.

## Acknowledgement

This work is supported by the National Institutes of Health grants R01GM108799 (to A. S. E. and D. F. C.), R01GM108580 (to G.A.), T32GM108539 (to A. S. E. and W. W. C.) and K08GM126336 (to W. W. C.); an International Anesthesia Research Society (IARS) mentored research award (to W. W. C.); The National Science Foundation grant DGE-1143954 (to M. M. B.) and the Taylor Family Institute for Innovative Psychiatric Research.

## Contribution

Z.W.C, J.R.B., D.F.C., G.A, D.E.R and A.S.E wrote the manuscript; Z.W.C, W.W.C, G.A, D.F.C., and A.S.E contributed to the project design; Z.W.C. developed middle-down MS methodology; Z.W.C. and M.M.B prepared MS samples and performed high-resolution LC-MS and data analysis; J.R.B created and maintained high-density GABA_A_ receptor expression cell line; Z.W.C and B.D.M conducted protein chemistry; A.L.G, D.J.S, G.A. performed electrophysiology; K.K., M.X.Q, D.F.C synthesized photolabeling compounds and click reaction linker; M.M.B designed and synthesized *FLI* tag; D.E.R performed computation modeling simulation and reagent docking.

## Supplemental Materials

### Material and Method

#### cDNA constructs

The human α_1_ and β_3_ subunits were subcloned into pcDNA3 for molecular manipulations and cRNA synthesis. Using QuikChange mutagenesis (Angilent), a FLAG tag was first added to the αι subunit then an 8xHis tag was added to generate the following His-FLAG tag tandem (QPSLHHHHHHHHDYKDDDDKDEL), inserted between the 4^th^ and 5^th^ residues of the mature peptide. The α1 and β3 subunits were then transferred into the pcDNA4/TO and pcDNA5/TO vectors (ThermoFisher), respectively, for tetracycline inducible expression. For *Xenopus laevis* Oocytes, point mutations were generated using the QuikChange site-directed mutagenesis kit (Agilent Technologies, Santa Clara, CA) and the coding region fully sequenced prior to use. The cDNAs were linearized with Xba I (NEB Labs, Ipswich, MA), and the cRNAs were generated using T7 mMessage mMachine (Ambion, Austin, TX).

#### Cell culture

The tetracycline inducible cell line HEK T-Rex™-293 (ThermoFisher) was cultured under the following conditions: cells were maintained in DMEM/F-12 50/50 medium containing 10% fetal bovine serum (tetracycline-free, Takara, Mountain View, CA), penicillin (100 units/ml), streptomycin (100 g/ml), and blastcidine (2 μg/ml) in a humidified atmosphere containing 5% CO2. Cells were passaged twice each week, maintaining subconfluent cultures. Stably transfected cells were cultured as above with the addition of hygromycin (50 μg/ml) and Zeocin (20 μg/ml).

#### Generation of high expression stable cell line

A stable cell line was generated by transfecting HEK T-Rex™-293 cells with human α_1_-8x His-FLAG pcDNA4/TO and human β_3_ pcDNA5/TO, in a 150 mm culture dish, using the Effectene transfection reagent (Qiagen). Two days after transfection, selection of stably transfected cells was performed with hygromycin and zeocin until distinct colonies appeared (usually after two weeks). Medium was exchanged several times each week to maintain antibiotic selection.

Individual clones (about 65) were selected from the dish and transferred to 24-well plates for expansion of each clone selected. When the cells grew to a sufficient number, about 50% confluency, they were split into two other plates, one for a surface ELISA against the FLAG epitope and a second for protein assay, to normalize surface expression to cell number^1^. The best eight clones were selected for expansion into 150 mm dishes, followed by [^3^H]muscimol binding. Once the best expressing clone was determined, the highest expressing cells of that clone were selected through fluorescence activated cell sorting (FACS).

#### FACS

FACS was done against the FLAG epitope, using a phycoerythrin (PE) conjugated anti-FLAG antibody. Fluorescent activated cells (1ml containing about 10 million cells) were sorted on the AriaII cell sorter (Washington University Pathology Core), collecting 0.5% of the highest fluorescing cells in a culture tube containing complete medium. The cells were plated in a 35mm dish and expanded until a near confluent 150 mm dish was obtained. Cells were enriched for expression by FACS three times. A final FACS was performed to select individual cells into a 96 well plate, which resulted in only 10 colonies of cells. These colonies were expanded and assayed for [^3^H]muscimol binding; the highest expressing clone was used for experiments.

#### Induction of GABA_A_ receptor expression

Stably transfected cells were plated into 50 150mm dishes. After reaching 50% confluency, GABA receptors were expressed by inducing cells with 1 μg/ml of doxycycline with the addition of 5mM sodium butyrate. Cells were harvested after 48 to 72 hours after induction.

#### Membrane protein preparation

HEK cells, after tetracycline induction, grown to :70–80% confluency were washed with 10 mM sodium phosphate/proteinase inhibitors (Sigma, USA) 2 times and harvested with cell scrapers. The cells were washed with 10 mM sodium phosphate/proteinase inhibitors and collected by centrifugation at 21,000g at 4°C for 5 min. The cells were homogenized with a glass mortar Teflon pestle for 10 strokes on ice. The pellet containing the membrane proteins was collected after centrifugation at 34,000g at 4°C for 30 min and resuspended in a buffer containing 10 mM potassium phosphate and 100mM KCl. The protein concentration was determined with micro-BCA protein assay and stored at −80 °C.

#### [^3^H]muscimol binding

[^3^H]muscimol binding assays were performed using a previously described method with minor modification^2^. Briefly, HEK cell membranes proteins (50 μg/ml final concentration) were incubated with :1–2 nM [^3^H]muscimol (30 Ci/mmol; PerkinElmer Life Sciences), neurosteroid in different concentrations (1nM-10 μM), binding buffer (10 mM potassium phosphate, 100 mM KCl, pH 7.5), in a total volume of 1 ml. Assay tubes were incubated for 1 h at 4 °C in the dark. Nonspecific binding was determined by binding in the presence of 1 mM GABA. Membranes were collected on Whatman/GF-C glass filter paper using a Brandel cell harvester (Gaithersburg, MD). To determine the Bmax of [^3^H]muscimol binding, 100 μg/ml of proteins were incubated with 250 nM [^3^H]muscimol, with specific activity reduced to 2Ci/mmol, for 1 hour at 4°C in the dark. The membranes were collected on Whatman/GF-B glass filter papers using manifold. Radioactivity bound to the filters was measured by liquid scintillation spectrometry using BioSafe II (Research Products International Corporation). Each data point was determined in triplicate.

#### Photolabeling of α_1_β_3_ GABA_A_ receptor

For all the photolabeling experiments, :10–20 mg of HEK cell membrane proteins (about 300 pmol [^3^H]muscimol binding) were thawed and resuspended in buffer containing 10 mM potassium phosphate, 100 mM KCl (pH 7.5) at a final concentration of 1.25 mg/ml. For photolabeling site identification experiments, 15 μM neurosteroid photolabeling reagent was added to the membrane proteins and incubated on ice for 1 hour. For the photolabeling competition experiments, 3 μM neurosteroid photolabeling reagent in the presence of 30 μM allopregnanolone or the same volume of ethanol were added for incubation. The samples were then irradiated in a quartz cuvette for 5 min, by using a photoreactor emitting light at >320 nm^2^. The membrane proteins were then collected by centrifugation at 20,000g for 45 min.

#### Cycloaddition (click reaction) of *FLI-tag* to KK123-photolabeled proteins

10mg of KK123 or ZCM42 photolabeled HEK membrane proteins were solubilized in 1 ml 2% SDS/PBS and incubated at RT for 2 hours. The protein lysate was collected by centrifugation at 21000g for 30 min. *FLI* tag was clicked to the KK123- or ZCM-photolabeled proteins at RT overnight in PBS buffer containing 2% SDS, 100 μM FLI tag^3^, 2.5mM sodium ascorbate, 250 μM Tris [(1-benzyl-1H-1,2,3triazol-4-yl)methyl]amine, 2.5 mM CuSO_4_. 1% Triton/PBS was added to the protein lysate to a SDS final concentration. of 0.05%. The protein lysate was loaded onto a streptavidin agarose column. The flow through was reloaded to the column two times or till the flow through was colorless and the streptavidin column was dark orange yellow. The column was washed with 10ml 0.05% Triton/PBS and eluted by 10ml 100 mM sodium dithionite/0.05%Triton/PBS. The column was turned into colorless after elutions. The eluted proteins were concentrated into 100 microul with 30 kDa cutoff Ccentricon apparatus. The supernatant of the Centricon tube was added into SDS-sample loading buffer, loaded to a 10% SDS-PAGE and transferred to a PVDF membrane, followed by Western blot with anti-α1 or antiβ3 antibody.

#### Purification of α_1_β_3_ GABA_A_ receptors

The photolabeled membrane proteins were resuspended in lysis buffer containing 1% n-dodecyl-β-D-maltoside (DDM), 0.25% cholesteryl hemisuccinate (CHS), 50 mM Tris (pH 7.5), 150 mM NaCl, 2 mM CaCl_2_, 5 mM KCl, 5 mM MgCl_2_, 1 mM EDTA, 10% glycerol at a final concentration of 1mg/ml. The membrane protein suspension was homogenized using a Teflon pestle in a motor-driven homogenizer and incubated at 4 °C overnight. The protein lysate was centrifuged at 20,000g for 45 min and supernatant was incubated with 0.5 ml anti-FLAG agarose (Sigma) at 4 °C for 2 hours. The anti-FLAG agarose was then transferred to an empty column, followed by washing with 20 ml washing buffer (50mM triethylammonium bicarbonate and 0.05% DDM). The GABA_A_ receptors were eluted with ten 1ml 200 μg/ml FLAG peptide and 100 μg/ml 3XFLAG (ApexBio) in the washing buffer. The 10 ml effective elutions containing GABA_A_ receptors (tested by Western blot with anti-a1 or anti3 antibody) were concentrated by 100 kDa cutoff Centricon filters into 0.1 ml.

#### Middle-down mass spectrometry

The purified GABA_A_ receptors (100ul) were reduced by 5mM tris (2-carboxyethyl) phosphine (TCEP) at for 30 min followed by alkylation with 7.5 mM N-ethylmaleimide (NEM) for 1 hour in the dark. The NEM was quenched by 7.5mM dithiothreitol (DTT) for 15 min. These three steps were done at RT. 8 μg of trypsin was added to the protein samples and incubated at 4 °C for :7–10 days. The digest was terminated by adding formic acid (FA) in a final concentration of 1%. The samples were then analyzed by an OrbiTrap ELITE mass spectrometer (Thermo Fisher Scientific) as in previous work ^4,5^ with some modifications. Briefly, a 20 μl aliquot was injected by an autosampler (Eksigent) at a flow rate of 800 nl/min onto a home-packed polymeric reverse phase PLRP-S column (Angilent, 12 cm × 75 μm, 300 Å). An acetonitrile (ACN) :10–90% concentration gradient was applied in the flow rate of 800 nl/min for 145 min to separate peptides. Solvent A: 0.1% FA/water and Solvent B: 0.1%FA/ACN. The ACN gradient was as follows: Isocratic elution at 10% solvent B, :1–60 min; :10–90% solvent B, :60–125 min; 90% solvent B, :125–135 min; 90%-10% solvent B, :135–140 min; isocratic solvent B, :140–145 min. For the first 60 min, a built-in divert valve on the mass spectrometer was used to remove the hydrophilic contaminants from the mass spectrometer. The survey MS1 scans were acquired at acquired at high resolution (60,000 resolution) in the range of m/z=:100–2000 and the fragmentation spectra were acquired at 15,000 resolution. Data dependent acquisition of the top 20 MS1 precursors with exclusion of singly charged precursors was set for MS2 scans. Fragmentation was performed using collision-induced dissociation or high-energy dissociation with normalized energy of 35%. The data were acquired and reviewed with Xcalibur 2.2 (Thermo Fisher Scientific).

#### MS data processing and analysis

The LC-MS data was searched against a customized database containing the sequence of the GABAA receptor 8XHis-FLAG-α_1_ and β_3_ subunit and filtered with 1% false discovery rate using PEAKS 8.5 (Bioinformatics Solutions Inc.). Search parameters were set for a precursor mass accuracy of 30 ppm, fragmentation ion accuracy of 0.1 Da, up to three missed cleavage on either side of peptide with trypsin digestion. Methionine oxidation, cysteine alkylation with NEM and DTT, any amino acids with adduct of KK123 (mass=372.16), KK200 (mass=462.27), KK202 (mass=500.31), KK123 with light *FLI* tag (mass=672.4322), and KK123 with heavy *FLI* tag (mass=682.44) were included as variable modification.

#### Receptor expression in *Xenopus laevis* Oocytes

The GABA_A_ receptors were expressed in oocytes from the African clawed frog *(Xenopus laevis)*. Frogs were purchased from Xenopus 1 (Dexter, MI), and housed and cared for in a Washington University Animal Care Facility under the supervision of the Washington University Division of Comparative Medicine. Harvesting of oocytes was conducted under the Guide for the Care and Use of Laboratory Animals as adopted and promulgated by the National Institutes of Health. The animal protocol was approved by the Animal Studies Committee of Washington University in St. Louis (approval No. 20170071).

The oocytes were injected with a total of 12 ng cRNA in 5:1 ratio (α1:β3) to minimize the expression of β3 homomeric receptors. Following injection the oocytes were incubated in ND96 with supplements (96 mM NaCl, 2 mM KCl, 1.8 mM CaCl_2_, 1 mM MgCl_2_, 2.5 mM Na pyruvate, 5 mM HEPES, and 100 U/ml+100 μg/ml penicillin+streptomycin and 50 μg/ml gentamycin; pH 7.4) at 16 ^o^C for :1–2 days prior to conducting electrophysiological recordings.

#### Electrophysiological recording

The electrophysiological recordings were conducted using standard two-electrode voltage clamp. Borosilicate capillary glass tubing (G120F-4, 0D=1.20 mm, ID=0.69 mm, Warner Instruments, Hamden, CT) were used for voltage and current electrodes. The oocytes were clamped at −60 mV. The chamber (RC-1Z, Warner Instruments, Hamden, CT) was perfused with ND96 at :5–8 ml min^−1^. Solutions were gravity-applied from 30-ml glass syringes with glass luer slips via Teflon tubing.

The current responses were amplified with an OC-725C amplifier (Warner Instruments, Hamden, CT), digitized with a Digidata 1200 series digitizer (Molecular Devices), and stored using pClamp (Molecular Devices). The peak amplitude was determined using Clampfit (Molecular Devices).

The stock solution of GABA was made in ND96 bath solution at 500 mM, stored in aliquots at −20°C, and diluted as needed on the day of experiment. Stock solution of propofol (200 mM in DMSO) was stored at room temperature. The steroids were dissolved in DMSO at 10 mM and stored at room temperature.

#### Electrophysiology data analysis

The αιβ_3_ wild-type and mutant receptors were tested (see table 1 and supplemental table 2 and 3) for potentiation by steroids (3α5α-allopregnanolone, 3α5β-pregnanolone, KK123, KK200 and KK-202) and direct activation by steroids (allopregnanolone KK123, KK200 KK-202 and pregnanolone). As control, several receptor isoforms were tested for potentiation by propofol. For each receptor type, we also determined constitutive open probability (P_o,const_).

To estimate P_o,const_, the effect of 100 μΜ picrotoxin (estimated P_o_ = 0) on the holding current was compared to the peak response to saturating GABA + 100 μΜ propofol (estimated P_o_= 1). P_o,const_ was then calculated as I_picrotoxin_/ (I_picrotoxin_-I_GABA+propofol_)^6^

Potentiation is expressed as the potentiation response ratio, calculated as the ratio of the peak response to GABA + modulator (steroid or propofol) to the peak response to GABA alone. The concentration of GABA was selected to produce a response of :5–15% of the response to saturating GABA + 100 μM propofol.

Direct activation by steroids was evaluated by comparing the peak response to 10 μM neurosteroid to the peak response to saturating GABA + 100 μM propofol. Direct activation by steroids is expressed in units of open probability that includes constitutive open probability. All data are given as mean ± S.D.

#### Docking simulations

An homology model of the α_1_β_3_ GAB_A_A receptor was developed using the crystal structure of the human β_3_ homopentamer published in 2014 (PDB Id: 4COF)^7^. In this structure the large cytoplasmic loops were replaced with the sequence SQPARAA used by Jansen et al.^8^ The pentamer subunits were organized as A α_1_, B β_3_, C α_1_, D β_3_, E β_3_. The alpha1sequence was aligned to the β_3_ sequence using the program MUSCLE^9^. The pentameric alignment was then used as input for the program Modeller^10^, using 4COF as the template, a total of 25 models were generated. The best model as evaluated by the DOPE score^11^ was then oriented into a POPC membrane and the system was fully solvated with 40715 TIP3 water molecules and ionic strength set to 0.15M KCl. A 100 ns molecular dynamics trajectory was then obtained using the CHARMM36 force field and NAMD. The resulting trajectory was then processed using the utility mdtraj^12^, to extract a snapshot of the receptor at each nanosecond of time frame. These structures were then mutually aligned by fitting the alpha carbons, providing a set of 100 mutually aligned structures used for docking studies.

The docking was performed using AutoDock Vina^13^ on each of the 100 snapshots in order to capture the receptor flexibility. Docking boxes were built for the beta3 intra-subunit site (cluster 1), the alpha1 intrasubunit site (cluster 2) and the alpha1-beta3 intersubunit site (cluster 3). The boxes were centered around the residues photolabeled by KK123, KK200, and KK202 and had dimensions of 25 × 25 × 25 Ångstroms, large enough to easily fit the linear dimensions of all of the steroids. For docking studies of allopregnanolone the docking boxes were placed in the same locations but had smaller dimensions of 20 × 20 × 20 Ångstroms. Docking was limited to an energy range of 3 kcal from the best docking pose, and limited to a total of twenty unique poses. The docking results for a given site could result in a maximum of 2000 unique poses (20 poses Χ 100 receptor structures), these were then clustered geometrically using the program DIVCF^14^. The resulting clusters were then ranked by Vina score, cluster size, and visually analyzed for compatibility with the photolabeling results, which is the photolabeling group oriented in the correct direction to produce the observed photo adducts.

##### Chemicals

The inorganic salts used in the buffers, GABA, picrotoxin, and the steroids 3α, 5α-allopregnanolone and 3α,5β-pregnanolone were purchased from Sigma-Aldrich (St. Louis). Propofol was purchased from MP Biomedicals (Solon).

**Chemical syntheses**. Solvents were either used as purchased or dried and purified by standard methodology. Extraction solvents were dried with anhydrous Na_2_SO_4_ and after filtration, removed on a rotary evaporator. Flash column chromatography was performed using silica gel (:32–63 μm) purchased from Scientific Adsorbents (Atlanta, GA). NMR spectra were recorded in CDCl_3_ (unless stated otherwise) at ambient temperature at 400 MHz (^1^H) or 100 MHz (^13^C).

##### Synthesis of 3-[4-(iodomethyl)phenyl]-3-(trifluoromethyl)-3H-Diazirine (8)

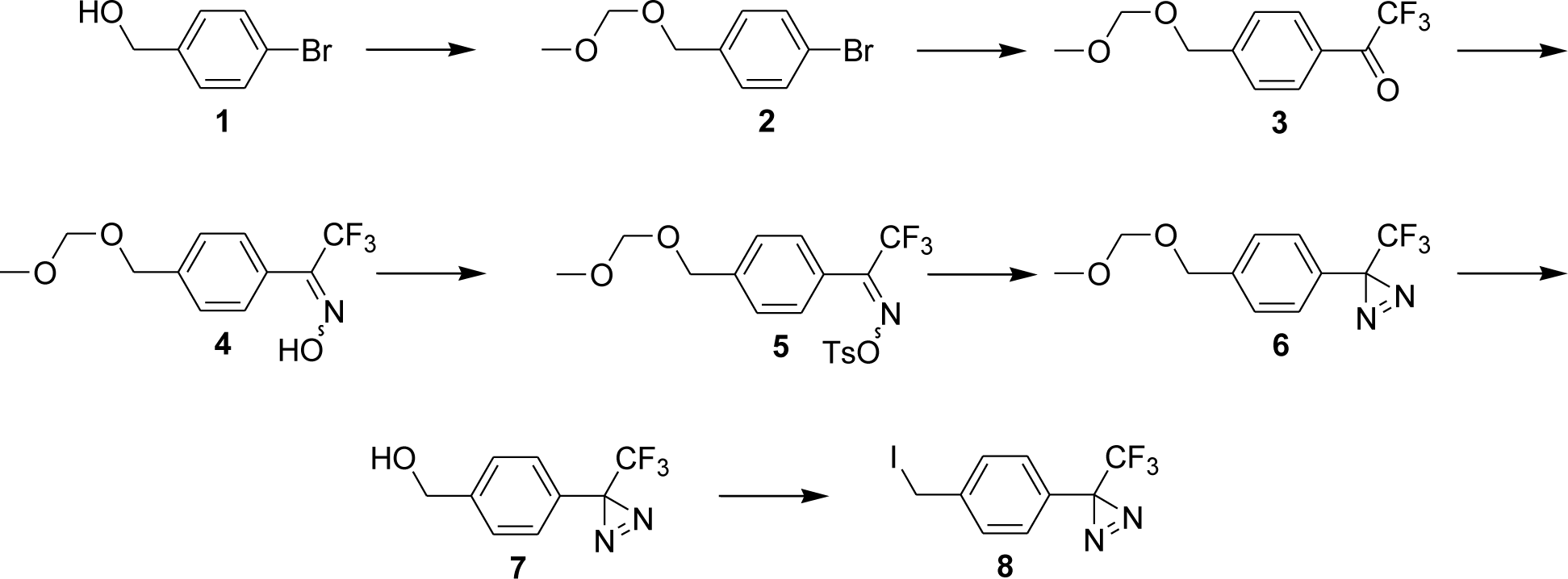

#### 1-Bromo-4-[(methoxymethoxy)methyl]-benzene (2)

Chloromethyl methyl ether (3.06 ml, 40.35 mmol) was added to a stirred, cold solution of 4-bromobenzyl alcohol (1, 5.0 g, 26.9 mmol) and Hunig’s base (14.05 ml, 80.7 mmol) in CH_2_Cl_2_ (30 ml) and the reaction was stirred at room temperature for 12 h. Aqueous NaHCO_3_ (100 ml) was added and the product was extracted into CH_2_Cl_2_ (3 × 75 ml). The combined extracts were dried, filtered and the solvents removed to give an oil which was purified by flash column chromatography (silica gel eluted with :5–10% EtOAc in hexanes) to give compound **2** as a colorless liquid (6.0 g, 97%) which had: ^1^H NMR δ 7.39, (d, 2H, *J* = 8.2 Hz), 7.16 (d, 2H, *J* = 8.2 Hz), 4.61 (s, 2H), 4.46 (s, 2H), 3.32 (s, 3H); ^13^C NMR δ 136.91, 131.49, 129.43, 121.52, 95.70, 68.36, 55.38.

#### 1-[4-[(Methoxymethoxy)methyl]phenyl]-2,2,2-trifluoro-ethanone (3)

Compound **2** (6.0 g, 26 mmol) in THF was added to a stirred hot suspension of Mg turnings (947 mg, 39 mmol) in THF (100 ml) and the mixture was heated to reflux for 15 min. Ethyl bromide (0.15 ml, 2 mmol) was added to the refluxing solution to activate the magnesium and the mixture was refluxed for another 90 min. The resulting Grignard reagent in THF was cooled to room temperature and transferred using a cannula into a flask containing a cold solution of 1-trifluoroacetyl piperidine (18.1 g, 100 mmol) in THF (15 ml). The reaction was allowed to stir for another 13 h at room temperature. Saturated aqueous NH_4_Cl was added and the product was extracted thrice into EtOAc. The combined extracts were dried, filtered and the solvents removed to give a pale yellow oil which was purified by flash column chromatography (silica gel eluted initially hexanes and then with :2–8% EtOAc in hexanes) to give compound **3** (4 g, 62%) as a liquid which had: ^1^H NMR δ 8.07 (d, 2H, *J* = 7.8 Hz), 7.54 (d, 2H, *J* = 7.8 Hz), 4.75 (s, 2H), 4.70 (s, 2H), 3.42 (s, 3H).

#### 1-[4-[(Methoxymethoxy)methyl]phenyl]-2,2,2-trifluoro-ethanone oxime (4)

Compound **3** (3.8 g, 15 mmol), hydroxylamine hydrochloride (6.95 g, 100 mmol) and NaOAc (16.4 g, 200 mmol) in MeOH (200 ml) were refluxed for 48 h. The reaction was cooled and the MeOH was removed. Water was added and the product was extracted trice into CH_2_Cl_2_. The combined extracts were dried, filtered and the solvents removed to give an *E/Z* mixture of compound **4** as an oil (3.8 g, 96%) which had: ^1^H NMR *(E/Z* mixture) δ 9.28 & 9.05 (b s, 1H), 7.:20–7.50 (m, 4H), 4.76 & 4.74 (s, 2H), 4.66 & 4.65 (s, 2H), 3.43(s, 3H).

#### 1-[4-[(Methoxymethoxy)methyl]phenyl]-2,2,2-trifluoro-ethanone,0-[(4-methylphenyl)sulfonyl] oxime (5)

Tosyl chloride (3.29 g 17.3 mmol) was added to a stirred, cold solution of compound **4** (3.8 g, 14.4 mmol) and triethyl amine (4.2 ml, 30 mmol) in CH_2_Cl_2_ and the reaction was stirred at 0 °C for 2 h. Aqueous saturated NaHCO_3_ was added and the product was extracted thrice into CH_2_Cl_2_. The combined extracts were dried, filtered and the solvents removed to give a residue which was then converted to compound 6 without purification. Compound **5** had: ^1^H NMR *(E/Z* mixture) δ 7.90 (m, 2H), 7.:30–7.55 (m, 6H), 4.75 & 4.72 (s, 2H), 4.65 & 4.64 (s, 2H), 3.43 & 3.42 (s, 3H), 2.49 & 2.47 (s, 3H).

#### 3-[4-[(Methoxymethoxy)methyl]phenyl]-3-(trifluoromethyl)-3H-diazirine (6)

Freshly condensed anhydrous ammonia (20 mL) was added to a stirred, cold (−78 °C) solution of compound **5** (6.0 g, 14.4 mmol) in CH_2_Cl_2_ (100 ml) and the reaction was slowly warmed to room temperature and stirred for 16 h. Water was added and the product was extracted into CH_2_Cl_2_ (3 × 80 ml). The combined extracts were dried, filtered and the solvent removed to give the crude diaziridine product which was dissolved in MeOH (50 ml). Triethyl amine (10 ml) was added and then MeOH saturated with I_2_ was added until the I_2_ color persisted. 5% Aqueous sodium thiosulfate (50 ml) and then water (100 ml) were added. The product was extracted into CH2Cl2 (3 × 75 ml). The combined extracts were dried, filtered and the solvents removed to give an oil which was purified by flash column chromatography (silica gel eluted with :2–10% EtOAc in hexanes) to give compound **6** (2.6 g, 70%) as a liquid which had: ^1^H NMR δ 7.40 (d, 2H, *J* = 8.2 Hz), 7.19 (d, 2H, *J* = 8.2 Hz), 4.71 (s, 2H), 4.61 (s, 2H), 3.41(s, 3H).

#### 4-[3-(Trifluoromethyl)-3H-diazirin-3-yl]-benzenemethanol (7)

Compound 6 (2.6 g, 10 mmol) in MeOH (10 ml) was added to :5–7 % dry HCl in MeOH (20 ml) and the reaction was stirred at room temperature for 13 h. Aqueous saturated NaHCO_3_ (100 ml) was added and the product was extracted into CH_2_Cl_2_ (3 × 75 ml). The combined extracts were dried, filtered and the solvent removed to give an oil which was purified by flash column chromatography (silica gel eluted with

5.20% EtOAc in hexanes) to give compound 7 (2.1 g, 97%) which had: ^1^H NMR δ 7.39 (d, 2H, *J* = 7.8 Hz), 7.20 (d, 2H, *J* = 7.8 Hz), 4.71 (s, 2H); ^13^C NMR δ 142.42, 128.15, 126.94, 126.45, 122.08 (q, *J* = 275 Hz), 63.90, 28.25 (q, *J* = 40.4 Hz).

#### 3-[4-(Iodomethyl)phenyl]-3-(trifluoromethyl)-3H-diazirine (8)

Imidazole (0.82 g, 12 mmol), triphenylphosphine (1.83g, 7 mmol) and I_2_ (2.03g, 8 mmol) in CH_2_Cl_2_ (20 ml) were stirred at room temperature for 15 min. Compound 7 (864 mg, 4 mmol) in CH_2_Cl_2_ (10 ml) was added and the reaction was stirred for 90 min at room temperature. Hexanes (100 ml) were added and the biphasic solution was stirred for 10 min. The supernatant hexanes, which were brownish in color, were added to a to a column of flash column silica gel and eluted with :2–10% EtOAc in hexanes to give compound **8** (1.1 g, 85%) as a viscous liquid which on standing crystallized to give a pale yellow solid. Compound **8** had: ^1^H NMR δ 7.40 (d, 2H, *J* = 7.8 Hz), 7.12 (d, 2H, *J* = 7.8 Hz), 4.43 (s, 2H); ^13^C NMR δ 141.07, 129.11 (2 x C), 128.65, 126.88 (2 x C), 121.99 (q, *J* = 275 Hz), 28.31 (q, *J* = 40.5 Hz).

##### Synthesis of KK-200 (12)

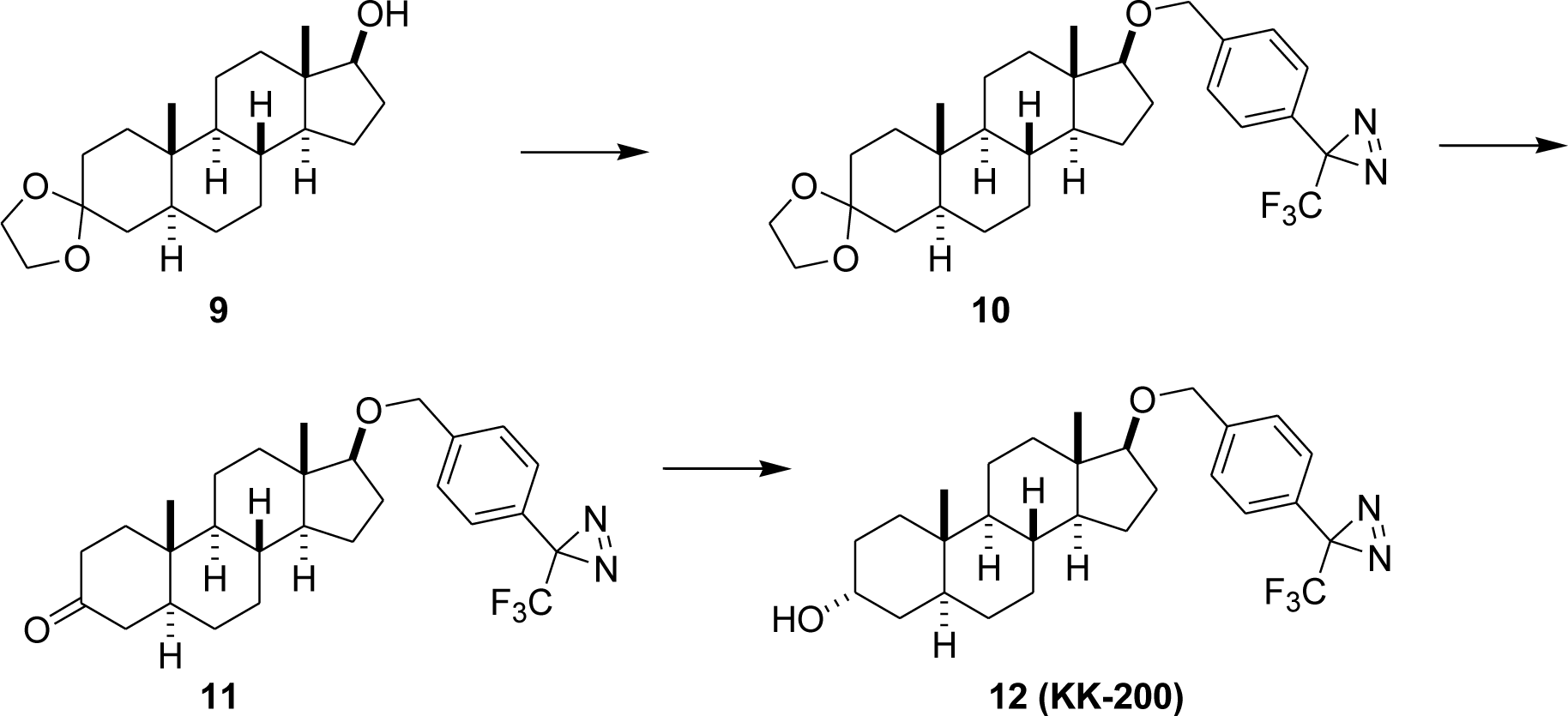

#### Steroid 10

Steroid **9** (301 mg, 0.9 mmol) and NaH (60% suspension in mineral oil, 0.8 g, 20 mmol) in DMF (10 ml) and THF (7 ml) were stirred at room temperature for 30 min. Compound **8** (652 mg, 2 mmol) was added and the reaction was stirred for 14 h. The reaction was cooled and 2-propanol was carefully added followed by the addition of cold water. When no NaH remained, ice followed by water (100 ml) was added and the product was extracted into EtOAc (4 × ml). The combined extracts were washed with brine, dried, filtered and the solvents removed to give an oil which was purified by flash column chromatography (silica gel eluted with hexanes followed by :2–20% EtOAc in hexanes) to give steroid **10** as an oil (205 mg, 43%) which had: ^1^H NMR: δ 7.36 (d, 2H, *J* = 8.2 Hz), 7.15 (d, 2H, *J* = 8.2 Hz), 4.53 (s, 2H), 3.92 (s, 4H), 3.38 (t, 1H, *J* = 8.2 Hz), 2.:05–0.60 (m), 0.82 (s, 3H), 0.81 (s, 3H); ^13^C NMR δ 141.27, 127.83, 127.37 (2 × C), 126.34 (2 × C), 122.10 (q, *J* = 275 Hz), 109.28, 88.68, 70.72, 64.07, 54.11, 51.13, 43.64, 43.12, 37.95, 37.91, 35.98, 35.49, 35.22, 31.54, 31.40, 31.08, 28.52, 28.37, 28.12, 27.84, 23.33, 22.61, 20.79, 14.07, 11.82, 11.35.

Steroid 9 has been prepared previously^15^.

#### Steroid 11

Steroid **10** (197 mg, 0.37 mmol) and *p*-toluenesulfonic acid (50 mg) in acetone (25 ml) were stirred at room temperature for 16 h. Aqueous saturated NaHCO3 was added and the acetone was removed. Water was added and the product was extracted into EtOAc (3 × 50 ml). The combined extracts were washed with brine, dried and the solvents removed. The crude product was purified by flash column chromatography (silica gel eluted with :15–20% EtOAc in hexanes) to give steroid **11** as an oil (163 mg, 90%) which had: ^1^H NMR: δ 7.36 (d, 2H, *J* = 8.2 Hz), 7.16 (d, 2H, *J* = 8.2 Hz), 4.53 (s, 2H), 3.38 (t, 1H, *J* = 8.2 Hz), 2.:45–0.60 (m), 1.01 (s, 3H), 0.83 (s, 3H); ^13^C NMR: δ 212.03, 141.19, 127.92, 127.43 (2 × C), 126.39 (2 × C), 122.12 (q, J = 275 Hz), 88.56, 70.76, 53.88, 50.95, 46.68, 44.66, 43.12, 38.53, 38.12, 37.79, 35.70, 35.16, 31.21, 28.76, 27.83, 23.37, 21.05, 11.84, 11.45.

#### Steroid 12 (KK-200)

K-selectride (1 M in THF, 1 ml, 1 mmol) was added to steroid **11** (160 mg, 0.33 mmol) dissolved in THF at −78 °C and the reaction was stirred at −78 °C for 1h. Water (a few drops) was added and the reaction was brought to 0 °C. A 1:1 mixture of 50% aqueous H_2_O_2_ (5 ml) and aqueous 4 N NaOH (5 ml) was added and the reaction was stirred at room temperature for 90 min. Water was added and the product was extracted into EtOAc (3 × 50 ml). The combined extracts were washed with brine, dried, filtered and the solvents removed to give an oil which was purified by flash column chromatography (silica gel eluted with :20–40% EtOAc in hexanes) to give steroid **12 (KK-200)** as a white solid (140 mg, 87%): ^1^H NMR: δ 7.36 (d, 2H, *J* = 8.2 Hz), 7.16 (d, 2H, *J* = 8.2 Hz), 4.53 (s, 2H), 4.04 (s, 1H), 3.38 (t, 1H, *J* = 8.2 Hz), 2.:05–0.70 (m), 0.81 (s, 3H), 0.78 (s, 3H); ^13^C NMR: δ 141.28, 127.86, 127.41 (2 × C), 126.36 (2 × C), 122.12 (q, *J* = 275 Hz), 88.73, 70.75, 66.47, 54.42, 51.22, 43.12, 39.10, 37.97, 36.12, 35.83, 35.25, 32.16, 31.51, 28.96, 28.55, 28.38, 28.14, 27.84, 23.31, 20.38, 11.84, 11.15.

#### Synthesis of KK-202 (17)

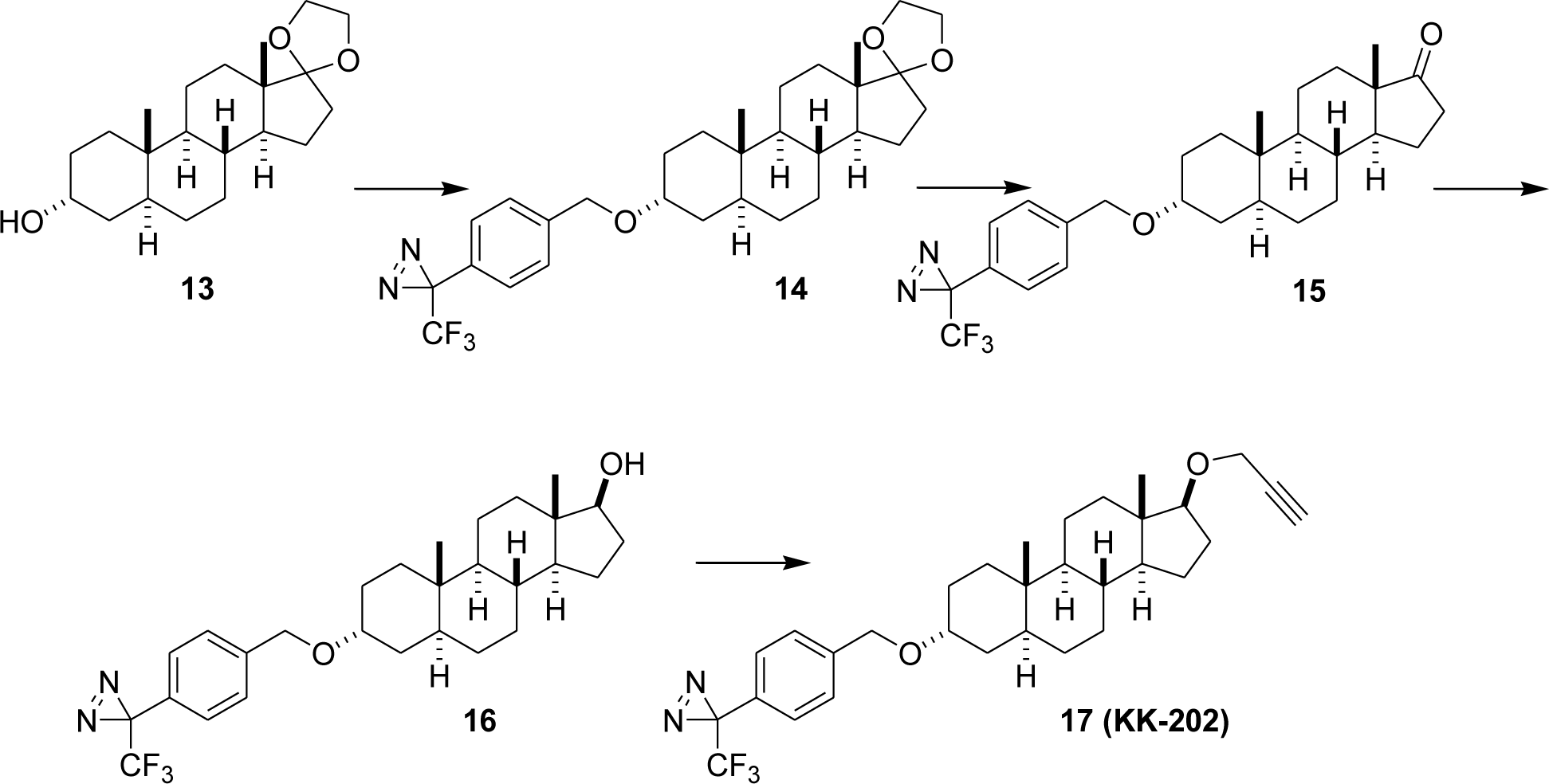

### Steroid 13

Commercially available androsterone (870 mg, 3.0 mmol), ethylene glycol (1 ml) and *p*-toluenesulfonic acid (200 mg) were refluxed for 15 h in a Dean Stark apparatus. The reaction was cooled and saturated aqueous NaHCO3 was added. The organic layer was washed with brine, dried, filtered and the solvents removed. The product was purified by flash column chromatography (silica gel eluted with :20–40% EtOAc in hexanes to give known steroid **13** (950 mg, 95%) as a solid which had: ^1^H NMR δ 4.02 (b s, 1H), 3.89 (m, 4H), 2.0 −0.75 (m), 0.82 (s, 3H), 0.77 (s, 3H); ^13^C NMR δ 119.45, 66.43, 65.10, 64.48, 54.02, 50.30, 45.91, 39.06, 36.08, 35.84, 35.70, 34.13, 32.15, 31.21, 30.66, 28.93, 28.38, 22.58, 20.12,14.37, 11.12.

Steroid **13** has been prepared previously from androsterone by the same procedure^16^.

### Steroid 14

Steroid **13** (301 mg, 0.9 mmol) and NaH (60% suspension in mineral oil, 0.8 g, 20 mmol) in DMF (10 ml) and THF (7 ml) were stirred at room temperature for 30 min. Compound **8** (652 mg, 2 mmol) was added and the reaction was stirred for 14 h. The reaction was cooled and 2-propanol was carefully added followed by cold water. When no NaH remained, ice was added to the mixture followed by water (100 ml) and the product was extracted into EtOAc (4 × 60m). The combined extracts were washed with brine, dried and the solvents removed to give an oil which was purified by flash column chromatography (silica gel eluted with hexanes followed by :2–20% EtOAc in hexanes) to give steroid **14** (240 mg, 50%) as an oil which had: ^1^H NMR δ 7.37 (d, 2H, *J* = 7.0 Hz), 7.18 (d, 2H, *J* = 7.0 Hz), 4.48 (s, 2H), 3.84 (m, 4H), 3.61 (s, 1H), 2.:01–0.80 (m), 0.84 (s, 3H), 0.79 (s, 3H); ^13^C NMR δ 141.40, 128.48, 127.50, 126.44, 122.14 (q, *J* = 275 Hz), 119.47, 73.6, 68.78, 65.13, 64.49, 54.08, 50.36, 45.96, 39.60, 35.94, 35.73, 34.15, 33.02, 32.78, 31.22, 30.69, 28.44, 25.63, 22.60, 22.00, 20.18, 14.38, 11.37.

### Steroid 15

Steroid **14** (230 mg, 0.43 mmol) and *p*-toluenesulfonic acid (100 mg) in acetone were stirred at room temperature for 15 h. Aqueous saturated NaHCO_3_ was added and the acetone was removed. The residue was dissolved in EtOAc and washed with brine, dried, filtered and the solvent removed to give an oil which was purified by flash column chromatography (silica gel eluted with hexanes followed by :5–25% EtOAc in hexanes) to give steroid **15** (195 mg, 93%) as an oil which had: ^1^H NMR δ 7.36 (d, 2H, *J* = 7.0 Hz), 7.16 (d, 2H, *J* = 7.0 Hz), 4.46 (AB q, 2H, *J* = 11.0 Hz), 3.60 (s, 1H), 2.:50–0.70 (m), 0.83 (s, 3H), 0.80 (s, 3H); ^13^C NMR δ 221.45, 141.40, 128.47, 127.53, 126.44, 122.13 (q, *J* = 275 Hz), 73.61, 68.88, 64.49, 54.35, 51.46, 47.79, 39.60, 36.05, 35.81, 35.01, 33.04, 32.69, 31.54, 30.75, 28.25, 25.53, 22.01, 21.71, 20.03, 13.77, 11.37.

### Steroid 16

NaBH_4_ (20 mg, 0.5 mmol) was added to steroid **16** (80 mg, 0.16 mmol) dissolvend in EtOH (7 ml) and the reaction was stirred at room temperature for 4 h. Water was added and the product was extracted into EtOAc (3 × 40 ml). The combined extracts were washed with brine, dried, filtered and the solvents removed to give an oil which was purified by flash column chromatography (silica gel eluted with hexanes followed by :2–20% EtOAc in hexanes) to give steroid **16** (60 mg, 74%) as an oil which had: ^1^H NMR δ 7.39 (d, 2H, *J* = 7.0 Hz), 7.18 (d, 2H, *J* = 7.0 Hz), 4.49 (AB q, 2H, *J* = 11.0 Hz), 3.63 (m, 2H), 2.:18–0.70 (m), 0.81 (s, 3H), 0.74 (s, 3H); ^13^C NMR δ 141.39, 128.30, 127.54, 126.44, 122.13 (q, *J* = 275 Hz), 81.96, 73.71, 68.85, 54.38, 51.05, 42.97, 39.67, 36.73, 36.00, 35.52, 33.05, 32.78, 31.49, 30.49, 28.45, 28.16, 25.62, 23.33, 20.35, 14.10, 11.69, 11.43, 11.12.

### Steroid 17 (KK-202)

Steroid **16** (40 mg, 0.08 mmol) and NaH (60% suspension in mineral oil, 80 mg, 2 mmol) in DMF (4 ml) were stirred at room temperature for 30 min. Propargyl bromide (80% w/v in toluene, 2 ml) was added and the reaction was stirred for 3 h during which time the reaction turned dark in color. The reaction was cooled to 0 °C and 2-propanol was carefully added followed by cold water. When no NaH remained, ice and then water (40 ml) were added and the product was extracted into EtOAc (4 × 30 ml). The combined extracts were washed with brine, dried, filtered and the solvents removed to give an oil which was purified by flash column chromatography (silica gel eluted with hexanes followed by :2–15% EtOH in hexanes to give crude steroid **17** (40 mg) as an oil which was purified in portions by preparative thin layer chromatography on three preparative plates that were twice eluted with 2% EtOAc in hexanes. Steroid **17** was visualized on the plate by brief exposure to iodine. Steroid 17 (KK-202) was an oil (16 mg, 37%) which had: ^1^H NMR δ 7.38 (d, 2H, *J* = 7.4 Hz), 7.17 (d, 2H, *J* = 7.4 Hz), 4.48 (AB q, 2H, *J* = 11.0Hz), 4.15 (m, 2H), 3.61 (s, 1H), 3.52 (t, *J* = 8.2 Hz), 2.39 (s, 1H), 2.:15–0.70 (m), 0.79 (s, 3H), 0.76 (s, 3H); ^13^C NMR δ 141.41, 127.88, 127.55, 126.45, 122.12 (q, *J* = 275 Hz), 88.31, 80.62, 73.72, 73.67, 68.86, 57.18, 51.20, 42.91, 39.63, 35.97, 35.26, 36.00, 35.52, 33.05, 32.77, 31.47, 28.45, 27.63, 25.62, 23.31, 20.39, 11.73, 11.42.

#### Synthesis of MQ-112 (22)

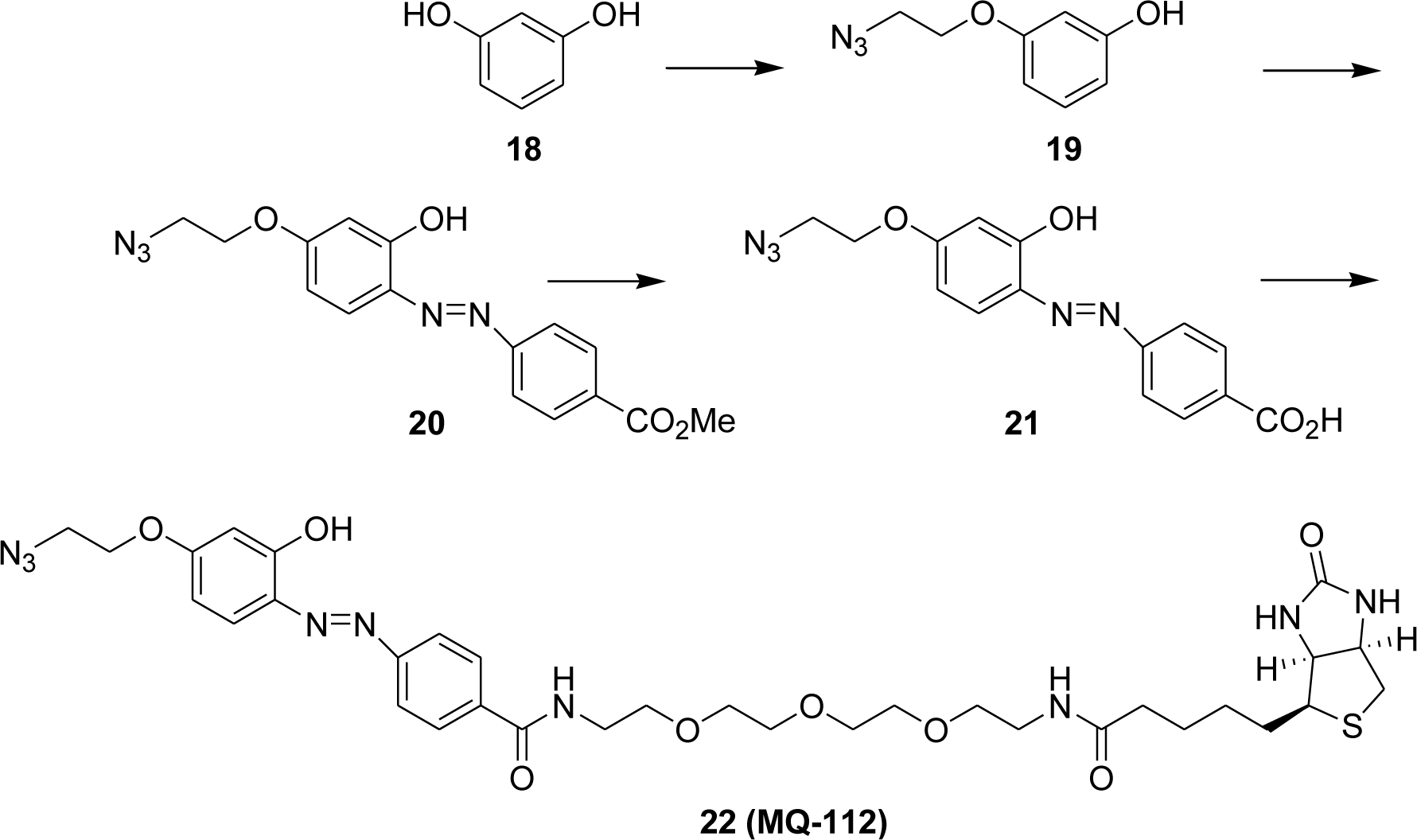

### Compound 19

Resorcinol (6 g, 54.4 mmol) and 2-azidoethyl 4-methylbenzenesulfonate (4.34 g, 18 mmol) were dissolved in EtOH (40 ml). KOH (1.14 g, 20.2 mmol, dissolved in 3.2 ml water) was slowly added and the reaction was heated to reflux for 3 h. After cooling to room temperature, the reaction was poured into water and the product extracted into Et_2_O (2 × 100 ml). The combined extracts were dried, filtered and the solvents removed to leave a residue which was purified by flash column chromatography (silica gel eluted with 10% EtOAc in hexanes) to give compound **19** (2.65 g, 82%) which had: ^1^H NMR δ 7.:13–7.10 (m, 1H), 6.:48–6.42 (m, 3H), 5.92 (s, br, 1H), 4.09 (t, *J* = 9.0 Hz, 2H), 3.55 (t, *J* = 9.0 Hz, 2H); ^13^C NMR δ 159.4, 156.8, 130.2, 108.5, 106.8, 102.2, 66.8, 50.0.

### Compound 20

NaNO_2_ (840 mg, 12 mmol) in H2O (24 ml) was added dropwise at 0 °C to a suspension of methyl 4-aminobenzoate (910 mg, 6 mmol) in 6 N HCl (24 ml). After 30 min, the clear solution was added dropwise to a solution of compound **19** (1.1 g, 6 mmol) in 2:1 H_2_O:THF (90 ml) containing K_2_CO_3_ (27 g). After addition was complete, 6 N HCl was added at 0 °C and the product was extracted into EtOAc (150 ml). The EtOAc was dried, filtered and the solvent removed to leave a residue which was purified by flash column chromatography (silica gel eluted with 10% EtOAc in hexanes) to give compound **20** (270 mg, 13%) which had: ^1^H NMR δ 13.86 (s, 1H), 8.14 (d, *J* = 8.7 Hz, 2H), 7.81 (d, *J* = 8.7 Hz, 2H), 7.74 (d, *J* = 8.6 Hz, 1H), 6.62 (d, *J* = 8.6 Hz, 1H), 6.42 (s, 1H), 4.17 (t, *J* = 9.0 Hz, 2H), 3.93 (s, 3H), 3.63 (t, *J* = 9.0 Hz, 2H); ^13^C NMR δ 166.3, 163.3, 157.9, 152.3, 135.2, 133.5, 130.7 (2 × C), 121.1 (2 × C), 109.4, 67.2, 52.2, 49.8.

### Compound 21

LiOH (0.3 g) dissolved in water (1 ml) was added to compound **20** (45 mg, 0.132 mmol) in THF/H_2_O (10 ml/10 ml). The mixture was stirred for 16 h, acidified with 6 N HCl until pH :4–5 and the product was extracted into EtOAc (100 ml). The combined extracts were washed with water (40 ml), dried, filtered and the solvents removed. The residue was purified by flash column chromatography (silica gel eluted with 10% MeOH in CH_2_Cl_2_) to give compound **21** (42 gm, 97%) which had: ^1^H NMR δ 8.11(d, *J* = 8.2 Hz, 2H), 7.96 (d, *J* = 8.2 Hz, 2H), 7.77 (d, 8.6 Hz, 1H), 6.:65–6.63 (m, 2H), 4.26 (t, *J* = 9.0 Hz, 2H), 3.63 (t, *J* = 9.0 Hz, 2H); ^13^C NMR δ 168.1, 163.6, 157.9, 153.5, 134.1, 132.0, 131.0 (2 × C), 127.7, 122.3 (2 × C), 109.0, 102.7, 67.9, 49.9.

Compound **21** has been prepared from resorcinol previously^17^.

### Compound 22

Anhydrous THF (8 ml) was added to an oven-dried flask under N_2_. Compound 21 (42 mg, 0.128 mmol), *N*-hydroxysuccinimide (49 mg, 0.33 mmol) and *N,N*’-dicyclohexylcarbodiimide (66 mg, 0.33 mmol) were added sequentially. The reaction was stirred at room temperature for 4 h. Solvent was removed under reduced pressure and the residue was dissolved partially in ice cold EtOAc. Undissolved material was removed by filtration and the filtrate was evaporated to dryness under high vacuum. The product, a NHS ester, was purified by flash column chromatography (silica gel eluted with 40% EtOAc in hexanes) and was dissolved in DMF (10 ml). Biotin-peg-NH_2_ (66 mg, 0.16 mmol) was added at room temperature. After 16 h, DMF was removed under high vacuum and the residue was purified by flash column chromatography (silica gel eluted with :5–10% MeOH in CH_2_Cl_2_) to give product (75 gm, 81%) which had: ^1^H NMR (CD3OD/CDCl_3_) δ 8.12 (d, *J* = 8.6 Hz, 2H), 8.01 (d, *J* = 8.2 Hz, 2H), 7.94 (d, *J* = 9.0 Hz, 1H), 6.82 (d, *J* = 2.0 Hz, 1H), 6.80 (d, *J* = 2.0 Hz, 1H), 4.:61–4.59 (m, 1H), 4.:43–4.36 (m, 3H), 3.:80–3.74 (m, 14H), 3.:67–3.64 (m, 2H), 3.:50–3.46 (m, 4H), 3.:30–3.25 (m, 1H), 3.:05–3.00 (m, 1H), 2.:86–2.82 (m, 1H), 2.33 (t, *J* = 7.4 Hz, 2H), 1.:84–1.39 (m, 9H); ^13^C NMR (CD_3_OD/CDCl_3_) δ 175.5, 168.9, 164.5, 158.1,152.9, 136.1, 135.4, 129.4 (2 × C), 122.2 (2 × C), 109.9, 102.9, 71.3 (2 × C), 70.9, 70.8, 70.4, 68.4, 62.9, 61.1, 56.6, 50.8, 41.0, 40.8, 40.0, 36.5, 29.4, 29.1, 26.4; LC-MS Calcd for [C_33_H_45_N_9_O_8_S+H^+^]: 728.3. Found: 728.4.

## Supplemental table 1

**Supplemental Table 1.** Summary of the photolabeling efficiency of the photolabeling analogues (15 μM) in their sites. The area under the curve of selected ion chromatography of photolabeled peptides was compared to that of the non-photolabeled peptides.

|  | KK123 (%) | KK200 (%) | KK202 (%) |
| --- | --- | --- | --- |
| $\alpha_1$-TM4 intrasubunit site | 0.38±0.04 | 0.06±0.08 | 0 |
| $\beta_3$-TM4 intrasubunit site | 3.19±2.26 | 0 | 0.53±0.63 |
| $\beta_3$- $\alpha_1$ intersubunit site | 0 | 1.54±0.03 | 0.42±0.23 |

**Supplemental Table 2.** Potentiation of α_1_β_3_ receptors GABA_A_ receptors in *Xenopus laevis* oocytes. Potentiation is expressed as potentiation response ratio, calculated as the ratio of the peak responses in the presence of GABA and neurosteroids to the peak response in the presence of GABA alone. The GABA concentrations were selected to generate a response of :5–15% of the response to saturating GABA. Data are shown as mean ± S.D. (number of cells).

| Receptor | Potentialiation by<br>0.1 μM 3α5αP | Potentialiation by<br>1 μM 3α5αP | Potentialiation by<br>0.1 μM 3α5βP | Potentialiation by<br>1 μM 3α5βP |
| --- | --- | --- | --- | --- |
| $\alpha_1\beta_3$ wild-type | 4.8±1.4 (6) | 8.4±3.4 (6) | 3.7±1.7 (6) | 7.7±4.3 (6) |
| $\alpha_1^{F407A}\beta_3$ | 5.3±2.1 (6) | 9.0±4.4 (6) | 3.0±0.9 (5) | 5.1±1.6 (5) |
| $\alpha_1^{N408W/Y411F}\beta_3$ | 1.4±0.2 (6) | 3.5±1.6 (10) | - | - |
| $\alpha_1^{W412A}\beta_3$ | 3.2±0.7 (5) | 6.3±2.2 (5) | 4.3±1.1 (6) | 7.1±2.7 (6) |
| $\alpha_1^{W412L}\beta_3$ | 5.2±1.4 (5) | 9.5±3.1 (5) | 5.2±1.6 (5) | 10.8±5.8 (5) |
| $\alpha_1^{Y415A}\beta_3$ | 4.9±1.7 (5) | 9.7±4.2 (5) | 3.9±1.4 (5) | 6.5±2.3 (5) |
| $\alpha_1^{F289A}\beta_3$ | 2.8±0.5 (6) | 4.9±1.0 (6) | 3.0±0.7 (4) | 5.5±1.3 (4) |
| $\alpha_1^{V227W}\beta_3$ | 2.0±0.6 (5) | 2.9±1.0 (5) | 2.2±0.7 (6) | 3.1±0.9 (6) |
| $\alpha_1\beta_3^{F438A}$ | 3.3±0.7 (5) | 5.7±2.1 (5) | 4.6±1.3 (5) | 11.9±6.9 (5) |
| $\alpha_1\beta_3^{W443A}$ | 4.9±1.4 (4) | 9.9±3.3 (4) | 3.5±0.6 (5) | 7.8±3.0 (5) |
| $\alpha_1\beta_3^{W443L}$ | 4.5±1.6 (5) | 13.9±2.6 (5) | 5.5±2.2 (5) | 12.6±4.9 (5) |
| $\alpha_1\beta_3^{Y445A}$ | 3.9±2.1 (10) | 11.7±4.7 (10) | 2.5±0.7 (9) | 6.8±4.3 (9) |
| $\alpha_1\beta_3^{Y284F}$ | 4.1±2.5 (10) | 10.3±2.9 (10) | 2.3±1.0 (5) | 7.2±1.9 (5) |
| $\alpha_1\beta_3^{I222W}$ | 4.0±0.6 (5) | 7.0±1.7 (5) | 3.2±0.5 (4) | 5.3±1.7 (4) |

**Supplemental Table 3.** Direct activation of α_1_β_3_ GABA_A_ receptors in *Xenopus laevis* oocytes by 10 μΜ allopregnanolone (3α5αΡ) and pregnanolone (3α5βΡ). Direct activation is expressed as in units of open probability. Data are shown as mean ± S.D. (number of cells).

| Receptor | Direct activation by 10 μM 3α5αP | Direct activation by 10 μM 3α5βP |
| --- | --- | --- |
| $\alpha_1\beta_3$ wild-type | 0.034±0.012 (5) | 0.110±0.052 (4) |
| $\alpha_1^{F407A}\beta_3$ | 0.073±0.035 (5) | 0.036±0.020 (5) |
| $\alpha_1^{N408W/Y411F}\beta_3$ | - | - |
| $\alpha_1^{W412A}\beta_3$ | 0.081±0.042 (5) | 0.070±0.027 (5) |
| $\alpha_1^{W412L}\beta_3$ | 0.18±0.10 (8) | 0.41±0.17 (5) |
| $\alpha_1^{Y415A}\beta_3$ | 0.44±0.13 (7) | 0.28±0.09 (10) |
| $\alpha_1^{F289A}\beta_3$ | 0.089±0.014 (4) | 0.057±0.011 (4) |
| $\alpha_1^{V227W}\beta_3$ | 0.012±0.017 (9) | 0.016±0.009 (5) |
| $\alpha_1\beta_3^{F438A}$ | 0.095±0.123 (8) | 0.089±0.049 (5) |
| $\alpha_1\beta_3^{W443A}$ | 0.10±0.05 (5) | 0.071±0.031 (5) |
| $\alpha_1\beta_3^{W443L}$ | 0.017±0.011 (5) | 0.018±0.015 (5) |
| $\alpha_1\beta_3^{Y445A}$ | 0.75±0.26 (7) | 0.66±0.10 (5) |
| $\alpha_1\beta_3^{Y284F}$ | 0.11±0.07 (6) | 0.09±0.05 (4) |
| $\alpha_1\beta_3^{I222W}$ | 0.23±0.26 (9) | 0.25±0.18 (10) |

## Supplemental Figure 1

**Supplemental Figure 1.**
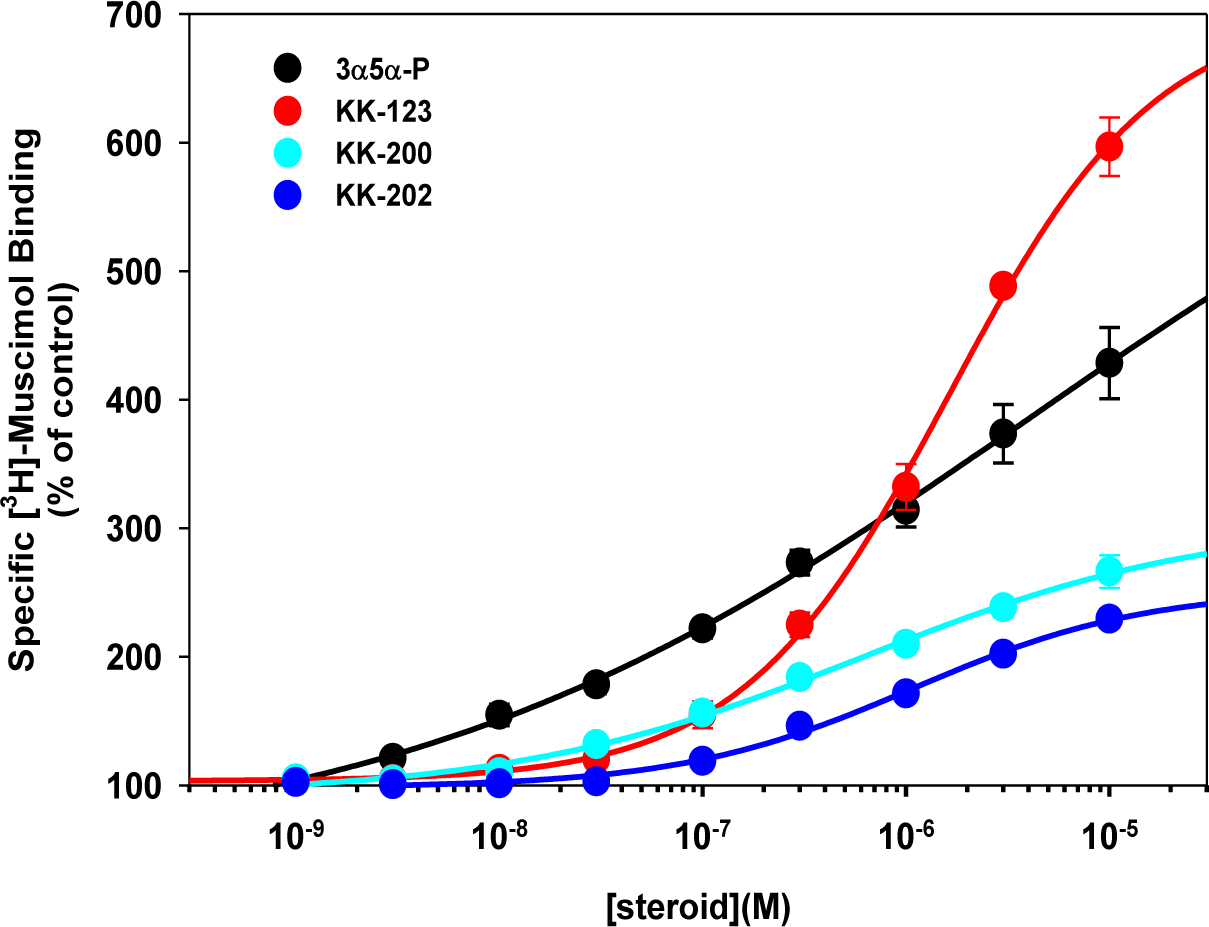
Enhancement of [^3^H]-muscimol binding of α_1_β_3_ GABA_A_ receptors by allopregnanolone and its photolabeling analogues. The EC_50_ (in mM) are 3.9±5.7 (n=9) for allopregnanolone; 1.6±0.2 (n=9) for KK123; 0.54±0.18 (n=9) for KK200; and 1.1±0.27 (n=9) for KK202.

**Supplemental Figure 2.**
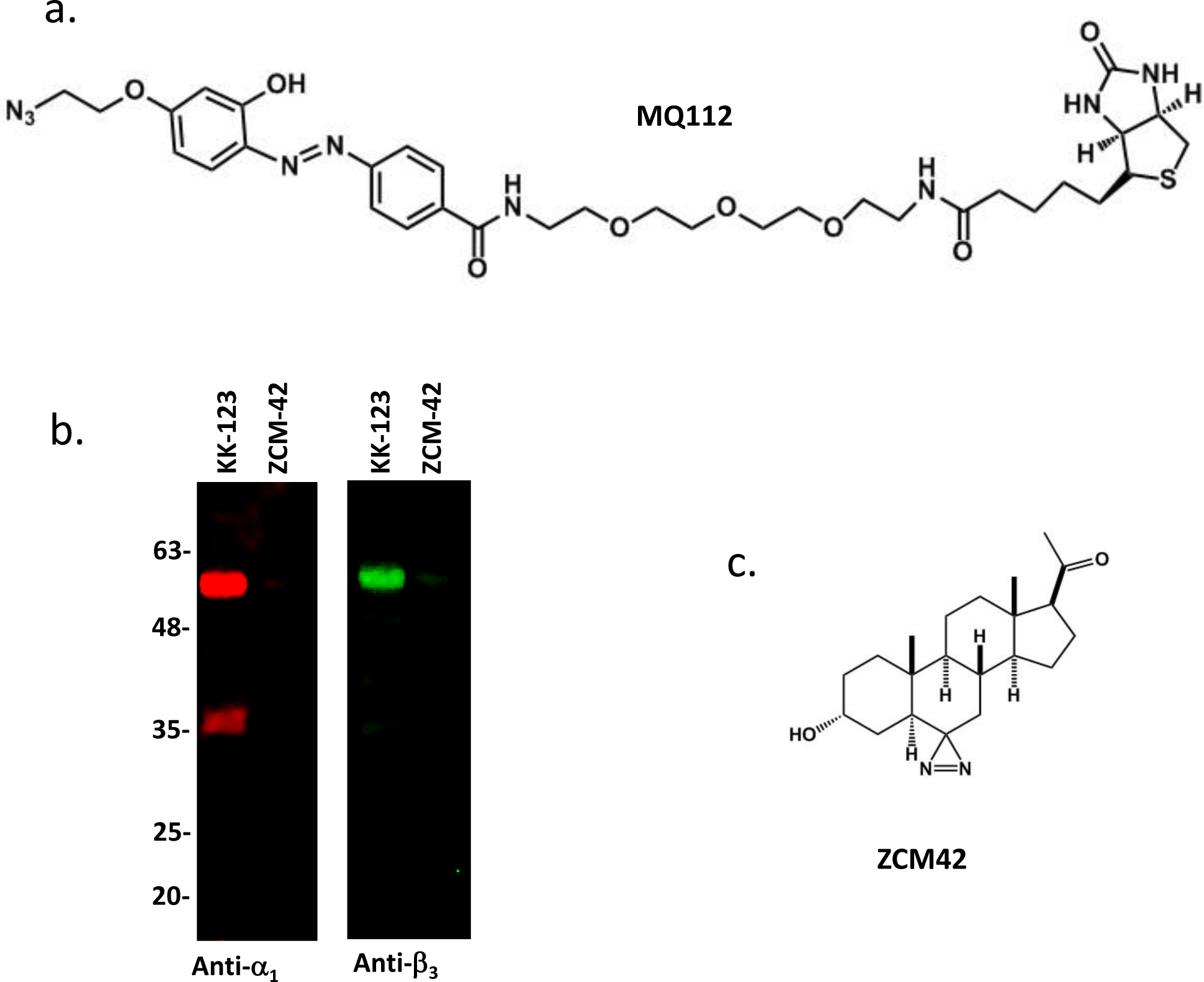
Purification of KK123 photolabeled GABA_A_ receptor via a trifunctional linker MQ112. a. The structure of MQ112. b. Purification of KK123 photolabeled GABA_A_ receptor α_1_ and β_3_ subunit by MQ112, via a click reaction, visualized by Western blot with anti-α_1_ and anti-β_3_. c. The structure of ZCM42.

**Supplemental Figure 3.**
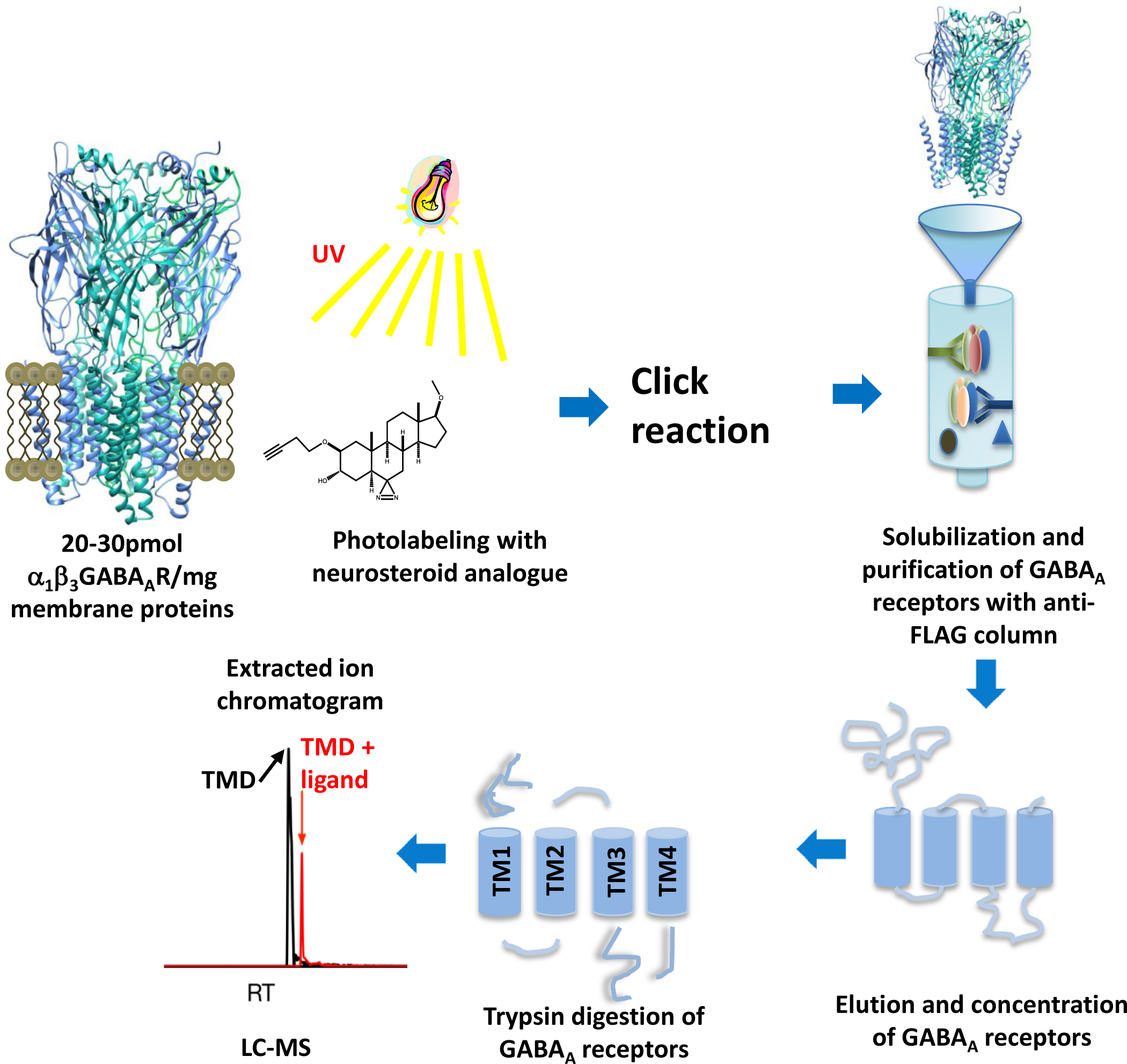
Workflow for identifying neurosteroid photolabeling sites in GABA_A_ receptors

**Supplemental Figure 4.**
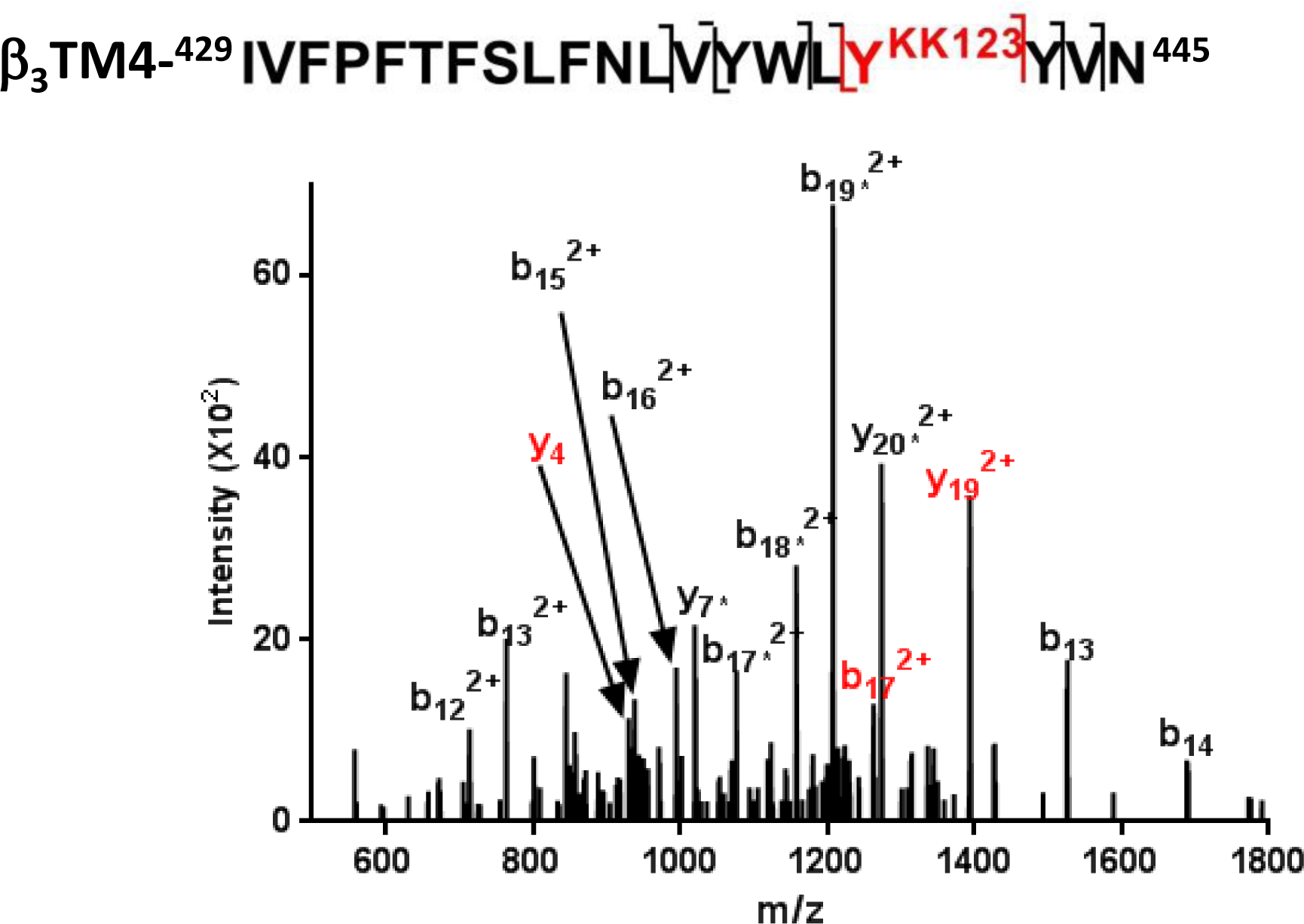
Fragmentation spectrum of KK123 photolabeled β_3_ΤΜ4 peptide. Y_4_ and b_17_ (in red) fragment ions containing a KK123 adduct indicate that Y^442^ is photolabeled by KK123. The fragment ions with neutral loss of the adduct are labeled as b_17_*^2^+, b_18_*^2^+, b_19_*^2+^, and y_20_*^2+^.

**Supplemental Figure 5.**
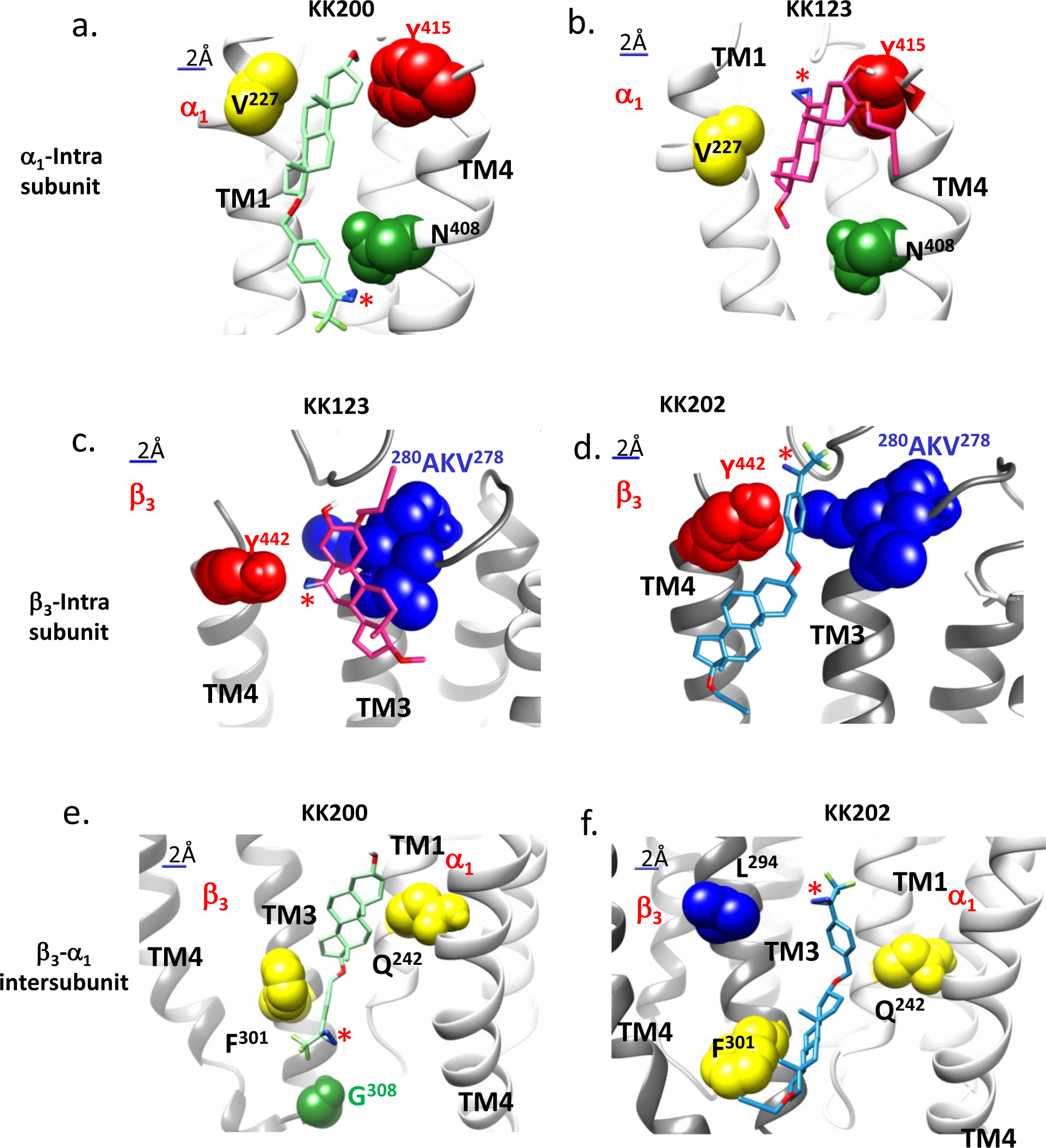
Computational docking of neurosteroid photolabeling analogues to their photolabeling sites on α_1_β_3_-GABA_A_ receptors. (a.) KK200 (light green) and (b.) KK123 (pink) in α_1_ intrasubunit site; (c.) KK123 and (d.) KK202 (light blue) in β_3_ intrasubunit site; (e.) KK200 and (f.) KK202 in β_3_ (+)/-α_1_(-) intersubunit site. The KK123 photolabeled residues α_1_-Y^415^ and β_3_-Y^442^ are colored red; the KK200 photolabeled residues α_1_-N^408^ and β_3_-G^308^ are colored green; and the KK202 β_3_-^278^VKA^280^ and L^294^ are colored blue. The canonical neurosteroid binding site Q^242^, F^301^, and the new mutation site V^227^ are colored yellow. The photolabeling diazirine group is indicated with red *.

## References

1. Agis-Balboa, R.C. et al. Characterization of brain neurons that express enzymes mediating neurosteroid biosynthesis. Proc Natl Acad Sci U S A 103, 14602–14607 (2006).

2. Faroni, A. & Magnaghi, V. The neurosteroid allopregnanolone modulates specific functions in central and peripheral glial cells. Front Endocrinol (Lausanne) 2, 103 (2011).

3. Baulieu, E.E. Neurosteroids: a novel function of the brain. Psychoneuroendocrinology 23, 963–987 (1998).

4. Barbaccia, M.L. et al. Endogenous γ-aminobutyric acid (GABA)A receptor active neurosteroids and the sedative/hypnotic action of γ-hydroxybutyric acid (GHB): A study in GHB-S (sensitive) and GHB-R (resistant) rat lines. Neuropharmacology 49, 48–58 (2005).

5. Kanes, S. et al. Brexanolone (SAGE-547 injection) in post-partum depression: a randomised controlled trial. Lancet 390, 480–489 (2017).

6. Reddy, D.S. & Rogawski, M.A. Enhanced anticonvulsant activity of ganaxolone after neurosteroid withdrawal in a rat model of catamenial epilepsy. J Pharmacol Exp Ther 294, 909–915 (2000).

7. Olsen, R.W. & Sieghart, W. GABA_A_ receptors: Subtypes provide diversity of function and pharmacology. Neuropharmacology 56, 141–148 (2009).

8. Gurley, D., Amin, J., Ross, P.C., Weiss, D.S. & White, G. Point mutations in the M2 region of the alpha, beta, or gamma subunit of the GABA_A_ channel that abolish block by picrotoxin. Receptors Channels 3, 13–20 (1995).

9. Yip, G.M. et al. A propofol binding site on mammalian GABA_A_ receptors identified by photolabeling. Nat Chem Biol 9, 715–720 (2013).

10. Jayakar, S.S. et al. Multiple propofol-binding sites in a gamma-aminobutyric acid type A receptor (GABAAR) identified using a photoreactive propofol analog. J Biol Chem 289, 27456–27468 (2014).

11. Li, G.D. et al. Identification of a GABA_A_ receptor anesthetic binding site at subunit interfaces by photolabeling with an etomidate analog. JNeurosci 26, 11599–11605 (2006).

12. Chiara, D.C. et al. Specificity of intersubunit general anesthetic-binding sites in the transmembrane domain of the human alpha1beta3gamma2 gamma-aminobutyric acid type A (GABAA) receptor. J Biol Chem 288, 19343–19357 (2013).

13. Chen, Z.W. et al. Neurosteroid analog photolabeling of a site in the third transmembrane domain of the beta3 subunit of the GABA(A) receptor. Mol Pharmacol 82, 408–419 (2012).

14. Hosie, A.M., Wilkins, ME., da Silva, H.M. & Smart, T.G. Endogenous neurosteroids regulate GABA_A_ receptors through two discrete transmembrane sites. Nature 444, 486–489 (2006).

15. Hosie, A.M., Wilkins, M.E. & Smart, T.G. Neurosteroid binding sites on GABA(A) receptors. Pharmacol Ther 116, 7–19 (2007).

16. Laverty, D. et al. Crystal structures of a GABA_A_-receptor chimera reveal new endogenous neurosteroid-binding sites. Nat Struct Mol Biol 24, 977–985 (2017).

17. Miller, P.S. et al. Structural basis for GABA_A_ receptor potentiation by neurosteroids. Nat Struct Mol Biol 24, 986–992 (2017).

18. Ziemba, A.M. et al. Alphaxalone Binds in Inner Transmembrane beta+-alpha-Interfaces of alpha1beta3gamma2 gamma-Aminobutyric Acid Type A Receptors. Anesthesiology 128, 338–351 (2018).

19. Akk, G. et al. The influence of the membrane on neurosteroid actions at GABA(A) receptors. Psychoneuroendocrinology 34 Suppl 1, S59–66 (2009).

20. Akk, G. et al. Neurosteroid access to the GABA_A_ receptor. J Neurosci 25, 11605–11613 (2005).

21. Chang, C.S., Olcese, R. & Olsen, R.W. A single M1 residue in the beta2 subunit alters channel gating of GABA_A_ receptor in anesthetic modulation and direct activation. J Biol Chem 278, 42821–42828 (2003).

22. Eaton, M.M. et al. Mutational Analysis of the Putative High-Affinity Propofol Binding Site in Human beta3 Homomeric GABA_A_ Receptors. Mol Pharmacol 88, 736–745 (2015).

23. Jurd, R. et al. General anesthetic actions in vivo strongly attenuated by a point mutation in the GABA(A) receptor beta3 subunit. FASEB J 17, 250–252 (2003).

24. Krasowski, M.D. et al. Propofol and other intravenous anesthetics have sites of action on the gamma-aminobutyric acid type A receptor distinct from that for isoflurane. Mol Pharmacol 53, 530–538 (1998).

25. Krasowski, M.D., Nishikawa, K., Nikolaeva, N., Lin, A. & Harrison, N.L. Methionine 286 in transmembrane domain 3 of the GABA_A_ receptor beta subunit controls a binding cavity for propofol and other alkylphenol general anesthetics. Neuropharmacology 41, 952–964 (2001).

26. Richardson, J.E. et al. A conserved tyrosine in the beta2 subunit M4 segment is a determinant of gamma-aminobutyric acid type A receptor sensitivity to propofol. Anesthesiology 107, 412–418 (2007).

27. Hosie, A.M., Clarke, L., da Silva, H. & Smart, T.G. Conserved site for neurosteroid modulation of GABA_A_ receptors. Neuropharmacology 56, 149–154 (2009).

28. Akk, G. et al. Neuroactive steroids have multiple actions to potentiate GABA_A_ receptors. J Physiol 558, 59–74 (2004).

29. Evers, A.S. et al. A synthetic 18-norsteroid distinguishes between two neuroactive steroid binding sites on GABA_A_ receptors. J Pharmacol Exp Ther 333, 404–413 (2010).

30. Li, P. et al. Natural and enantiomeric etiocholanolone interact with distinct sites on the rat alpha1beta2gamma2L GABA_A_ receptor. Mol Pharmacol 71, 1582–1590 (2007).

31. Cheng, W.W.L. et al. Mapping two neurosteroid-modulatory sites in the prototypic pentameric ligand-gated ion channel GLIC. J Biol Chem 293, 3013–3027 (2018).

32. Chiara, D.C. et al. Photoaffinity labeling the propofol binding site in GLIC. Biochemistry 53, 135–142 (2014).

33. Nury, H. et al. X-ray structures of general anaesthetics bound to a pentameric ligand-gated ion channel. Nature 469, 428–431 (2011).

34. Willenbring, D., Liu, L.T., Mowrey, D., Xu, Y. & Tang, P. Isoflurane alters the structure and dynamics of GLIC. Biophys J 101, 1905–1912 (2011).

35. Budelier, M.M. et al. Click Chemistry Reagent for Identification of Sites of Covalent Ligand Incorporation in Integral Membrane Proteins. Anal Chem 89, 2636–2644 (2017).

36. Barrantes, F.J. & Fantini, J. From hopanoids to cholesterol: Molecular clocks of pentameric ligand-gated ion channels. Progress in Lipid Research 63, 1–13 (2016).

37. Jiang, X. et al. A clickable neurosteroid photolabel reveals selective Golgi compartmentalization with preferential impact on proximal inhibition. Neuropharmacology 108, 193–206 (2016).

38. Das, J. Aliphatic diazirines as photoaffinity probes for proteins: recent developments. Chem Rev 111,:4405–4417 (2011).

39. Brunner, J. New Photolabeling and Crosslinking Methods. Annual Review of Biochemistry 62, 483–514 (1993).

40. Chen, Z.W. et al. 11-trifluoromethyl-phenyldiazirinyl neurosteroid analogues: potent general anesthetics and photolabeling reagents for GABA_A_ receptors. Psychopharmacology (Berl) 231, 3479–3491 (2014).

41. Chen, Z.W., Fuchs, K., Sieghart, W., Townsend, R.R. & Evers, A.S. Deep amino acid sequencing of native brain GABAA receptors using high-resolution mass spectrometry. Mol Cell Proteomics 11, M111 011445 (2012).

42. Shu, H.J. et al. Slow actions of neuroactive steroids at GABA_A_ receptors. J Neurosci 24, 6667–6675 (2004).

43. Miller, P.S. & Aricescu, A.R. Crystal structure of a human GABA_A_ receptor. Nature 512, 270–275 (2014).

44. Li, P., Bandyopadhyaya, A.K., Covey, D.F., Steinbach, J.H. & Akk, G. Hydrogen bonding between the 17beta-substituent of a neurosteroid and the GABA(A) receptor is not obligatory for channel potentiation. Br J Pharmacol 158, 1322–1329 (2009).

45. Unwin, N. Refined structure of the nicotinic acetylcholine receptor at 4A resolution. J Mol Biol 346, 967–989 (2005).

46. Mennerick, S. et al. Selective antagonism of 5alpha-reduced neurosteroid effects at GABA(A) receptors. Mol Pharmacol 65, 1191–1197 (2004).

## References

1 Bracamontes, J.R. & Steinbach, J.H. Multiple modes for conferring surface expression of homomeric beta1 GABA_A_ receptors. J Biol Chem 283, 26128–26136 (2008).

2 Darbandi-Tonkabon, R. et al. Photoaffinity labeling with a neuroactive steroid analogue. 6-azi-pregnanolone labels voltage-dependent anion channel-1 in rat brain. J Biol Chem 278, 13196–13206 (2003).

3 Budelier, M.M. et al. Click Chemistry Reagent for Identification of Sites of Covalent Ligand Incorporation in Integral Membrane Proteins. Anal Chem 89, 2636–2644 (2017).

4 Cheng, W.W.L. et al. Mapping two neurosteroid-modulatory sites in the prototypic pentameric ligand-gated ion channel GLIC. J Biol Chem 293, 3013–3027 (2018).

5 Chen, Z.W. et al. Neurosteroid analog photolabeling of a site in the third transmembrane domain of the beta3 subunit of the GABA(A) receptor. Mol Pharmacol 82, 408–419 (2012).

6 Eaton, M.M. et al. Multiple Non-Equivalent Interfaces Mediate Direct Activation of GABA_A_ Receptors by Propofol. Curr Neuropharmacol 14, 772–780 (2016).

7 Miller, P.S. & Aricescu, A.R. Crystal structure of a human GABA_A_ receptor. Nature 512, 270–275 (2014).

8 Jansen, M., Bali, M. & Akabas, M.H. Modular design of Cys-loop ligand-gated ion channels: functional 5-HT3 and GABA rho1 receptors lacking the large cytoplasmic M3M4 loop. J Gen Physiol 131, 137–146 (2008).

9 Edgar, R.C. MUSCLE: a multiple sequence alignment method with reduced time and space complexity. BMC Bioinformatics 5, 113 (2004).

10 Sali, A. & Blundell, T.L. Comparative protein modelling by satisfaction of spatial restraints. J Mol Biol 234, 779–815 (1993).

11 Shen, M.Y. & Sali, A. Statistical potential for assessment and prediction of protein structures. Protein Sci 15, 2507–2524 (2006).

12 McGibbon, R.T. et al. MDTraj: A Modern open library for the analysis of molecular dynamics Trajectories. Biophys J 109, 1528–1532 (2015).

13 Trott, O. & Olson, A.J. AutoDock Vina: improving the speed and accuracy of docking with a new scoring function, efficient optimization, and multithreading. JComput Chem 31, 455–461 (2010).

14 Meslamani, J.E., Andre, F. & Petitjean, M. Assessing the geometric diversity of cytochrome P450 ligand conformers by hierarchical clustering with a stop criterion. J Chem Inf Model 49, 330–337 (2009).

15 Mappus, E., Renaud, M., Rolland de Ravel M., Grenot, C. & Cuilleron, C.Y. Synthesis and characterization by ^1^H and ^13^C nuclear magnetic resonance spectroscopy of 17 alpha-hexanoic derivatives of 5 alpha-dihydrotestosterone and testosterone. Steroids 57, 122–134 (1992).

16 Tchedam Ngatcha, B., Luu-The, V., Labrie, F. & Poirier, D. Androsterone 3alpha-ether-3beta-substituted and androsterone 3beta-substituted derivatives as inhibitors of type 3 17beta-hydroxysteroid dehydrogenase: chemical synthesis and structure-activity relationship. J Med Chem 48, 5257–5268 (2005).

17 Yang, Y.Y., Grammel, M., Raghavan, A.S., Charron, G. & Hang, H.C. Comparative analysis of cleavable azobenzene-based affinity tags for bioorthogonal chemical proteomics. Chem Biol 17, 1212–1222 (2010).

